# EnzCast: Prediction of Patient-Specific Enzymatic Kinetics thrugh Multi-Modal Deep Learning and Isoform-Resolved Bayesian Inference based on Single-Cell Transcriptomics

**DOI:** 10.64898/2026.04.28.721430

**Authors:** Xuechen Mu, Yan Yang, Qingyu Wang, Zimin Chen, Bizhe Luo, Zhenyu Huang, Xinyi Lin, Long Xu, Xuan Li, Yinwei Qu, Jun Xiao, Zhihang Wang, Bocheng Shi, Qi Ou, Bowen Yao, Jing Yan, Yangmu Zhuang, Ye Zhang, Rui Shi, Ying Xu

## Abstract

Enzyme kinetic parameters underpin mechanistic biology but remain sparse in physiological context. We present EnzCast, a multi-modal framework jointly predicting *K*_m_, *k*_cat_, *k*_cat_/*K*_m_, and *K*_i_ from protein sequence, 3D structure, substrate chemistry, and experimental conditions, paired with IsoKin, an isoform-resolved Bayesian framework converting EnzCast priors into patient-specific *in vivo* kinetics. Trained on KinBench, the largest curated kinetics database, task-adaptive EnzCast achieved *R*^2^ = 0.413, 0.455, 0.227, and 0.105 for *K*_m_, *K*_i_, *k*_cat_, and *k*_cat_/*K*_m_, surpassing all baselines on catalytic tasks. Systematic condition scans recovered compartment-specific pH direction inversion and pathway-level temperature responses. In a 20-patient colorectal cancer single-cell cohort, IsoKin reduced posterior uncertainty by 73.3% and 77.3%, revealing cell-type-specific rewiring. Orthogonal validation—scFEA flux, DepMap essentiality (permutation *P* = 0.0008) and TCGA survival— provided mixed but directionally consistent support. Together, EnzCast and IsoKin bridge *in vitro* prediction, condition-aware biochemical interrogation and patient-resolved *in vivo* inference.

## 1 INTRODUCTION

Enzyme kinetic parameters—the Michaelis constant (*K*_m_), turnover number (*k*_cat_), catalytic efficiency (*k*_cat_/*K*_m_), and inhibition constant (*K*_i_)—are fundamental descriptors of catalytic functions that dictate the rates of metabolic fluxes, signaling strengths, and even the effectiveness of therapeutic strategies. Despite their centrality to quantitative biology, the availabilities of such data fall far short behind the levels of availabilities of omic data that have enabled large-scale systems biology studies, mainly because they remain predominantly determined through labor-intensive *in vitro* assays requiring purified enzymes and controlled conditions. As protein sequences and other omic data, such as transcriptomic and metabolomic data, have grown by orders of magnitude in the past two decades, the repositories for kinetic parameters of biochemical reactions have grown far more slowly, creating a progressively widening “kinetic gap” [1,2] that have greatly limited the utilities of the omic data that are being generated at explosive rates due to the rapid progress in the sequencing and other omic technologies. This has become the main bottleneck of realistic modeling of complex cellular behaviors, for which great amounts of omic data have been generated but remain substantially underutilized, such as in human diseases.

Measurements of kinetic data are inherently condition-dependent: pH, temperature, organism-dependent, and inhibition context can shift such kinetic parameters by orders of magnitude [3], and yet most available data reflect only the idealized laboratory conditions. This issue is particularly serious in human disease conditions, such as cancer or Alzheimer’s disease where the physicochemical conditions are profoundly altered from the physiological ones, including both the intracellular and extracellular pH [4,5], temperature [6,7], and the redox states [8]. On top of these, the levels of cofactor availabilities are altered, such as NAD^+^/NADH [9]. All these are known to affect the kinetic parameters, and hence the biochemical reaction pathways, the reaction consequence, and the cellular-level behaviors of the diseased cells under study. Yet no existing frameworks adequately bridge the gap between the needs for realistic mechanistic studies of human diseases and the lack of the adequate kinetic data needed for human disease studies.

Recent machine learning advances have enabled the development of deep learning models that can infer kinetic parameters directly from enzyme sequences and substrate structures. State-of-the-art methods such as DLKcat [10], UniKP [11], TurNuP [12], and CatPred [13] combine protein encoders—ranging from convolutional architectures to pretrained transformer-based protein language models—with molecular graph representations of substrates to predict *k*_cat_, *K*_m_, *K*_i_, or combinations thereof. These studies have shown that sequence- and structure-derived features contain information relevant to catalytic reactions and can generalize to unseen enzyme-substrate pairs. However, existing predictors face several fundamental limitations that restrict their utilities for patient-specific applications. First, they provide organism-agnostic, “average-condition” estimates that do not account for condition-dependent shifts in kinetic parameters. Second, architectural constraints limit context-aware predictions: most models encode enzyme and substrate in separate branches, fuse them via concatenation or shallow attention, and append environmental variables as terminal scalar features, such that the conditioning signal influences only the terminal layers rather than internal computations that map molecular representations to predictions. Third, models that treat each enzyme as a purely sequential object may struggle to capture local geometric determinants of catalysis arising from the three-dimensional configuration of residues forming the binding pocket and catalytic machinery. Fourth, *k*_cat_, *K*_m_, and *K*_i_ are related but not interchangeable, and the derived catalysis efficiency parameter *k*_cat_/*K*_m_is mathematically coupled to *k*_cat_and *K*_m_on the logarithmic scale; when these relationships are ignored or enforced too rigidly, models may either underutilize shared information or propagate inconsistencies. Most critically, existing models output point predictions without a principled mechanism to integrate patient-specific transcriptomic or proteomic data, leaving a conceptual gap between population-level kinetic estimates and the *in vivo* apparent parameters that shape metabolic fluxes in individual diseased cells.

Here we present EnzCast (Enzyme Kinetics Contextualized for Adaptive Stratification of Tumors), a framework that addresses these limitations through two connected advances. First, EnzCast is a multi-modal deep learning architecture that conditions kinetic parameter prediction on experimental context by integrating protein sequences (ESM-2 [14] with LoRA [15]), three-dimensional structures (equivariant graph neural networks [16]), substrate molecular graphs (graph isomorphism networks [17] with Morgan fingerprints [18]), and experimental conditions, such as pH, temperature, organism, enzyme type, reaction context, through learned adaptive scaling factors that modulate feature fusion. We trained EnzCast on KinBench, a curated benchmark of 359,652 kinetic measurements covering all four parameters from 25,118 enzymes and 23,547 substrates—4.7-fold larger than the largest prior benchmark (Extended Data Table 1) and the only benchmark simultaneously covering all four parameters, including *K*_i_ for inhibition prediction. Evaluated under stringent sequence-similarity clustering [19], EnzCast achieves task-adaptive accuracy with dedicated single-task experts for binding parameters and a multi-task model for catalytic parameters, outputting heteroscedastic uncertainty estimates as priors for downstream Bayesian inference. A dual-pathway prediction head groups *K*_m_/ *K*_i_ (binding) and *k*_cat_ / *k*_cat_/*K*_m_(catalytic) to leverage shared mechanistic information (Fig. 1, Modules 1–2). Systematic *in silico* condition scans further validate the condition encoder, showing compartment-specific pH direction inversion and pathway-level temperature responses consistent with biophysical principles.

**Fig. 1.**
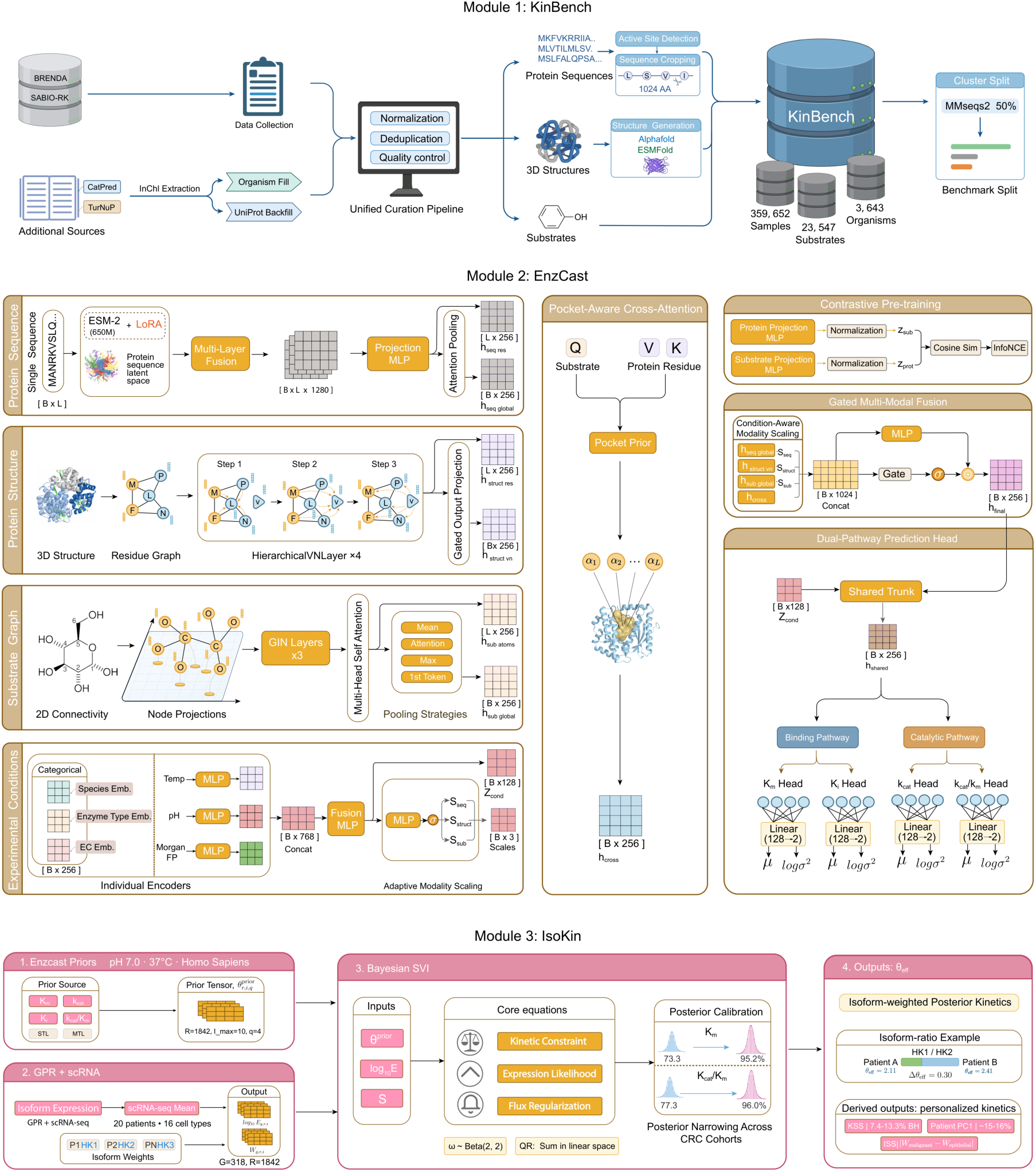
Overview of the EnzCast framework for patient-specific enzyme kinetics. Module 1 — KinBench. 359,652 records (SABIO-RK [22], BRENDA [23]): *K*_*m*_ (181,663), *k*_cat_ (99,173), *k*_cat_/*K*_*m*_ (39,799), *K*_*i*_ (39,017); 25,118 enzymes, 23,547 substrates, 3,643 organisms; MMseqs2 [19] (50%; 13,529 clusters) and random splits. Module 2 — EnzCast. ESM-2 [14] sequences, EGNN [16] structures, GIN [17] substrate graphs, and conditions encoded; binding (*K*_*m*_, *K*_*i*_) and catalytic (*k*_cat_, *k*_cat_/*K*_*m*_) outputs carry heteroscedastic uncertainty. Module 3 — IsoKin. EnzCast priors (pH 7.0, 37 °C) and scRNA-seq [25] via GPR decomposition, SVI [21] with Recon3D [20], and effective kinetics *θ*^eff^ = log_10_{∑_*i*_ *i*_*i*_ 10^*θ*^_*i*_|, yielding patient-specific parameters across 318 contexts.

Second, and central to our contribution, we introduce IsoKin, an isoform-resolved Bayesian calibration framework that integrates EnzCast’s tumor-contextualized kinetic priors generated under physiological intracellular conditions (pH 7.0, 37°C) with patient-specific single-cell RNA-seq data to infer personalized apparent *in vivo* kinetic parameters. Rather than regressing transcript abundance directly onto kinetic parameters, IsoKin expands the traditional “gene–protein–enzymatic reaction” line to include explicit isoforms, assigns isoform-level priors from EnzCast, and constrains posterior kinetic parameter prediction using a stoichiometric flux model [20] implemented with stochastic variational inference [21]. Context specificity emerges after inference by combining posterior isoform-based kinetic parameters with patient-and cell-type-specific fractions of isoform-level expressions, yielding effective reaction-level parameters for each cellular context.

Applied to a colorectal cancer single-cell cohort, the framework identifies patient- and cell-type-specific kinetic landscapes, detects malignant-versus-control metabolic rewiring, and exposes isoform-dependent effects that are inaccessible to gene-level analyses. Together with systematic condition scans and orthogonal analyses against scFEA flux, DepMap essentiality and TCGA survival, these results position EnzCast with IsoKin as a route from population-level kinetic prediction to biophysically grounded, patient-resolved metabolic hypotheses (Fig. 1, Module 3).

## 2 RESULTS

### 2.1 A unified benchmark for enzyme kinetic parameter estimation using KinBench

Systematic comparison of enzyme kinetic prediction methods has been hampered by the absence of a unified benchmark with standardized evaluation protocols. To address this, we constructed KinBench by integrating experimental kinetic measurements from two manually curated repositories—SABIO-RK [22] and BRENDA [23]—supplemented with data from published datasets generated using machine learning (ML) such as TurNuP [12] and CatPred [13]. The resulting benchmark covers four kinetic parameters: Michaelis constant (*K*_m_), turnover number (*k*_cat_), catalytic efficiency (*k*_cat_/*K*_m_), and inhibition constant (*K*_i_). In contrast, no existing benchmark covers all four kinetic parameters (Extended Data Table 1) as (1) DLKcat [10] and TurNuP [12] are *k*_cat_-focused benchmarks, (2) UniKP [11] extends the coverage to *K*_m_ and *k*_cat_/*K*_m_ but omits *K*_i_, and (3) CatPred [13] covers *k*_cat_, *K*_m_, and *K*_i_ but not *k*_cat_/*K*_m_.

All kinetic data were standardized to consistent units, log_10_-transformed, and subjected to canonical substrate normalization, UniProt [24] protein mapping, composite-key deduplication and active-site-centered sequence cropping for proteins exceeding 1,024 residues to preserve catalytically relevant spatial context for downstream structure encoding (METHODS).

KinBench currently comprises 359,652 records spanning 25,118 unique UniProt-mapped enzyme entries of 4,442 EC classes and 23,547 canonical substrates, together from 3,643 organisms (Fig. 2; Extended Data Table 1). *K*_m_entries constitute the largest subset (181,663; 50.5%), followed by *k*_cat_ (99,173; 27.6%), *k*_cat_/*K*_m_ (39,799; 11.1%), and *K*_i_ (39,017; 10.8%). It is noteworthy that the simultaneous inclusion of all four parameters enables to estimate their quantitative relationships, hence allowing to predict the missing values of one parameter, such as *k*_cat_/*K*_m_, which tends to be missing in the publicly available databases.

**Fig. 2.**
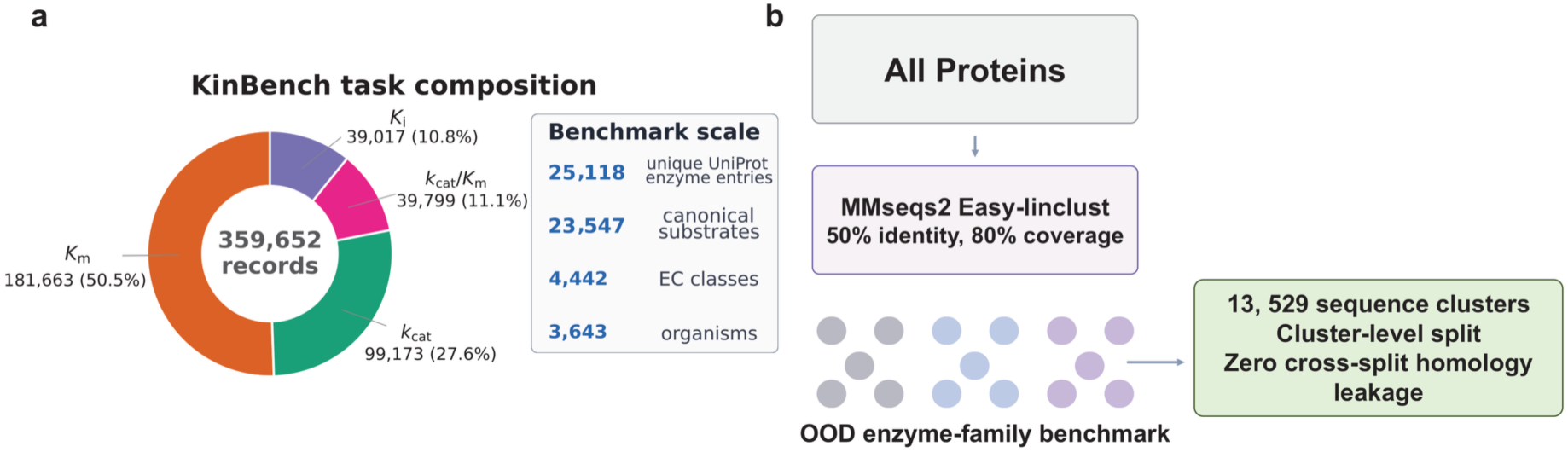
KinBench. A unified multi-task enzyme kinetics benchmark. a, Task composition of KinBench showing the distribution of 359,652 records across four kinetic parameters (*K*_*m*_, *k*_cat_, *k*_cat_/*K*_*m*_, *K*_*i*_) and scale comparison with existing benchmarks. b, MMseqs2-based sequence clustering and partitioning strategy: all unique protein sequences are clustered at 50% identity, and clusters are assigned to training, validation, and test sets to ensure zero sequence-homology overlap.

KinBench is 3.3–43.7× larger per task than the closest competitor (4.3× for *k*_cat_, 4.4× for *K*_m_, 43.7× for *k*_cat_/*K*_m_, and 3.3× for *K*_i_; Extended Data Table A1) and is the only benchmark applying a uniform curation workflow across all four parameters (Extended Data Fig. 1b–d) Even under condition-agnostic accounting that collapses replicate measurements to unique enzyme–substrate–organism triplets, as the existing benchmarks do not model experimental conditions (e.g., pH, temperature), KinBench retains 1.9–18.3× more entries per task than the closest competitor (Extended Data Fig. 1a); the remaining condition-specific records provide the training signal for EnzCast’s condition encoder, a capability absent from all the existing methods.

To evaluate generalization under realistic out-of-family conditions, we used sequence-similarity-based partitioning rather than random splitting, which can place close homologs in both training and test sets and inflate performance estimates. We clustered all unique protein sequences at 50% identity using MMseqs2 [19] (easy-linclust, 80% bidirectional coverage, cov-mode 0), yielding 13,529 non-redundant clusters, and partitioned data at the cluster level to minimize sequence-homology overlap among training, validation and test sets (Fig. 2). This design ensures that test-set enzymes belong to protein families absent from training data, directly probing extrapolation to evolutionarily distant enzyme families—the scenario most relevant to enzyme discovery and patient-specific applications.

All baselines were retrained on their original datasets to verify implementation fidelity before KinBench evaluation (Extended Data Table A5; Methods).

### 2.2 Reproducible, task-adaptive kinetic parameter predictions by EnzCast

We first trained EnzCast in multi-task mode across all four kinetic parameters and benchmarked it against four established methods—DLKcat [10], UniKP [11], TurNuP [12], and CatPred [13]—using matched KinBench UniRef50-clustered evaluation partitions ( < 50% sequence identity; Fig. 3). All baselines were evaluated with the same benchmark preprocessing and held-out test split (METHODS). Multi-task EnzCast achieved the best performance for the catalytic tasks: *R*^2^ = 0.227 for *k*_cat_ (*n* = 10,235 test samples) vs. UniKP ( *R*^2^ = 0.192), and *R*^2^ = 0.105 for *k*_cat_/*K*_m_ ( *n* = 4,063) vs. UniKP ( *R*^2^ = 0.072), with DLKcat yielding negative *R*^2^for both catalytic outputs and CatPred negative for *k*_cat_/*K*_m_(Fig. 3b,c). This performance advantage reflects the consistency constraint (Equation 3), which softly enforces the biochemical relationship log_10_(*k*_cat_/*K*_m_) = log_10_*k*_cat_ − log_10_*K*_m_ during joint training.

**Fig. 3.**
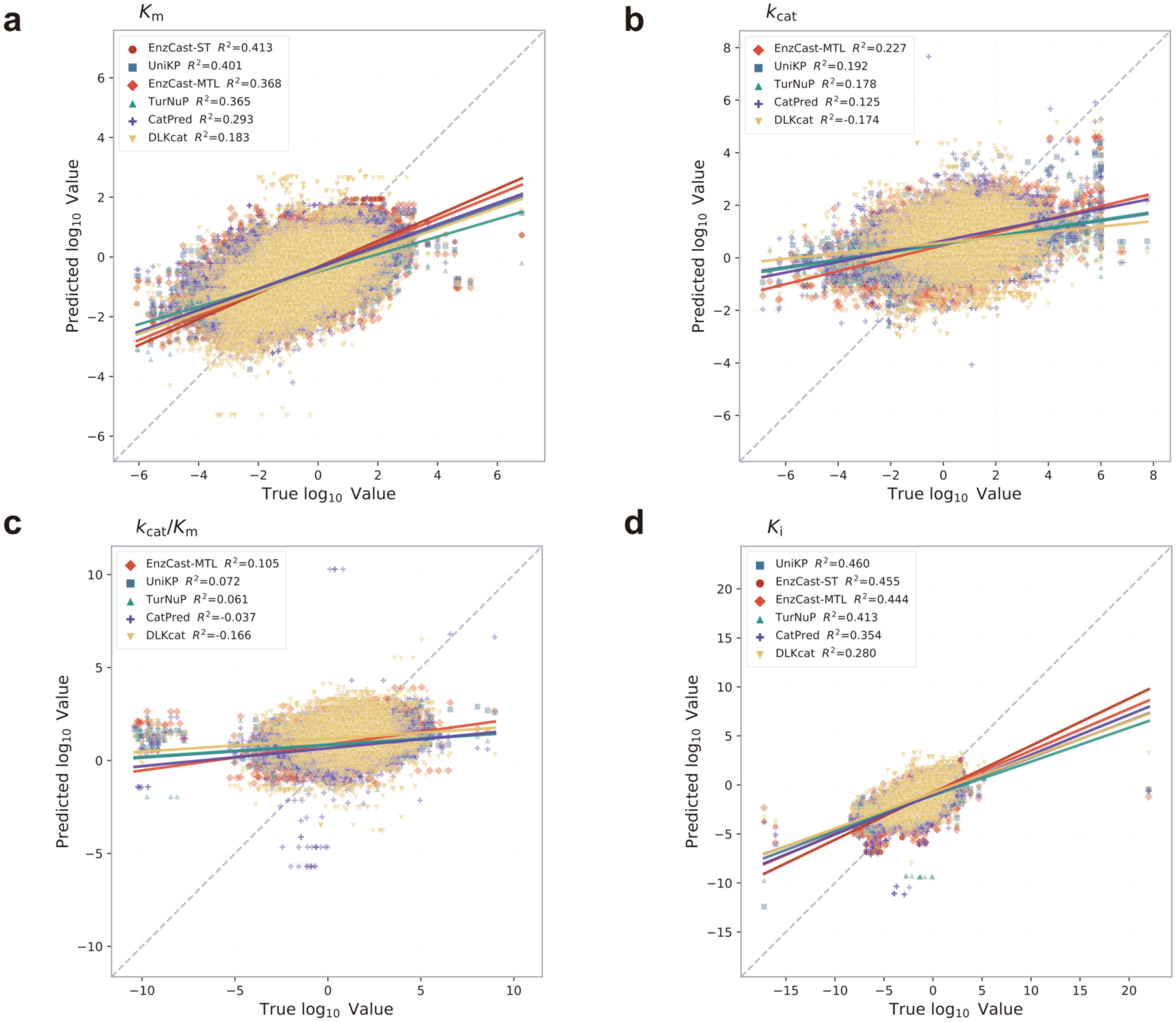
EnzCast yields reproducible, task-adaptive kinetic parameter predictions. a, Scatter of true versus predicted log_10_*K*_*m*_values for EnzCast-ST and four baselines on the KinBench UniRef50-clustered test set; legend entries report per-panel *R*^2^of the displayed representative checkpoint, sorted in descending order. EnzCast-ST achieves *R*^2^ = 0.413 (representative) and 0.404 ± 0.001 (multi-seed headline). b, Scatter for *k*_cat_using EnzCast-MTL (*R*^2^ = 0.227 representative; 0.234 ± 0.020 multi-seed headline). DLKcat yields negative *R*^2^. c, Scatter for *k*_cat_/*K*_*m*_using EnzCast-MTL ( *R*^2^ = 0.105 representative; 0.083 ± 0.007 multi-seed headline). d, Scatter for *K*_*i*_using EnzCast-ST ( *R*^2^ = 0.455 representative; 0.468 ± 0.013 multi-seed headline); in this panel, EnzCast-ST and UniKP are closely matched, whereas the multi-seed summary exceeds the baseline. Multi-seed reproducibility and MTL versus ST comparison are shown in Extended Data Fig. 2.

For the binding tasks, however, multi-task EnzCast fell slightly below the best baselines: *R*^2^ = 0.368 vs. UniKP 0.401 for *K*_m_ ( *n* = 18,915) and *R*^2^ = 0.444 vs. UniKP 0.460 for *K*_i_ ( *n* = 3,777). This performance by the multi-task predictors motivated dedicated single-task training for *K*_m_ and *K*_i_, giving rise to improved performance by single-task EnzCast at *R*^2^ = 0.413 for *K*_m_, exceeding UniKP’s 0.401, and *R*^2^ = 0.455 for *K*_i_, approaching UniKP’s 0.460 (Fig. 3a,d). Conversely, single-task training reduced *k*_cat_/*K*_m_ performance from *R*^2^ = 0.105 to 0.022 (Extended Data Table 2), confirming that the consistency constraint is essential for catalytic efficiency.

We therefore adopted a task-adaptive benchmark configuration: multi-task learning (EnzCast-MTL) for *k*_cat_and *k*_cat_/*K*_m_and single-task training (EnzCast-ST) for *K*_m_and *K*_i_(METHODS). Using this configuration, EnzCast exceeded the best baseline predictor for Km and both catalytic tasks in the displayed benchmark, while closely approaching UniKP for *K*_i_ and clearly surpassing DLKcat with *R*^2^ = 0.183 for *K*_m_ and negative values for both catalytic outputs, and CatPred with *R*^2^ = 0.293 for *K*_m_ and a negative value for *k*_cat_/*K*_m_. The generally lower *R*^2^ values across all methods compared with their original publications reflect the substantially larger scale and greater enzymatic diversity of KinBench.

Repeated training across three random seeds (see METHODS) confirmed the high reproducibility of EnzCast predictions of *K*_m_and *K*_i_ with mean *R*^2^ = 0.404 ± 0.001 for *K*_m_ (coefficient of variation (CV) 0.15%) and 0.468 ± 0.013 for *K*_i_ (CV 2.67%), while predictions for the catalytic parameters had larger but bounded variability, with mean *R*^2^ = 0.234 ± 0.020 for *k*_cat_ (CV 8.47%) and 0.083 ± 0.007 for *k*_cat_/*K*_m_(CV 8.45%) (Extended Data Fig. 2a).

Systematic ablation analyses of the 3D structure encoder, cross-attention module, experimental-condition encoder, *β*-NLL loss, and dual-pathway prediction head showed that removing any single element led to reduced performance in at least one task, with cross-attention producing the largest average drop across the task-adaptive benchmark and the dual-pathway head acting as a second broad contributor (Extended Data Table 3; Extended Data Fig. 2c). Each prediction also retained a heteroscedastic variance term, providing the uncertainty-aware priors that the downstream Bayesian calibration framework requires.

### 2.3 Condition-aware predictions capture biophysically consistent kinetic responses

We performed systematic condition scans across 1,561 human enzymes in KinBench to test whether EnzCast’s condition encoder learned biophysically meaningful responses rather than condition-specific memorization (METHODS). For pH, we exploited the reversed pH gradient in tumor cells—intracellular alkalinization (pH 7.0 → 7.4) and extracellular acidification (pH 7.4 → 6.5)—and classified enzymes by subcellular localization (589 cytoplasmic, 166 secreted/extracellular) and theoretical isoelectric point (pI). Under intracellular alkalinization, catalytic efficiency (*k*_cat_/*K*_m_) increased consistently (Cohen’s *d* = 0.29 − 0.54; Fig. 4a,b), whereas extracellular acidification produced the opposite response with larger effect sizes (*d* = −0.97 to − 1.28; Fig. 4d; per-pI-bin statistics in Extended Data Fig. 12). Critically, for 166 secreted/extracellular enzymes evaluated under both pH regimes, intracellular and extracellular *k*_cat_ shifts were strongly anti-correlated (Spearman *ρ* = −0.70, *P* = 8.0 × 10^−26^; Fig. 4c), demonstrating that the same enzyme set exhibits opposite kinetic responses under opposite compartmental pH shifts—a direction inversion learned from training data alone, without an explicit biophysical prior. We did not detect a significant *k*_cat_gradient across pI bins (Kruskal–Wallis *P* = 0.38), so the supported claim is compartment-level pH directionality rather than fine-grained pI-ranked responsiveness.

**Fig. 4.**
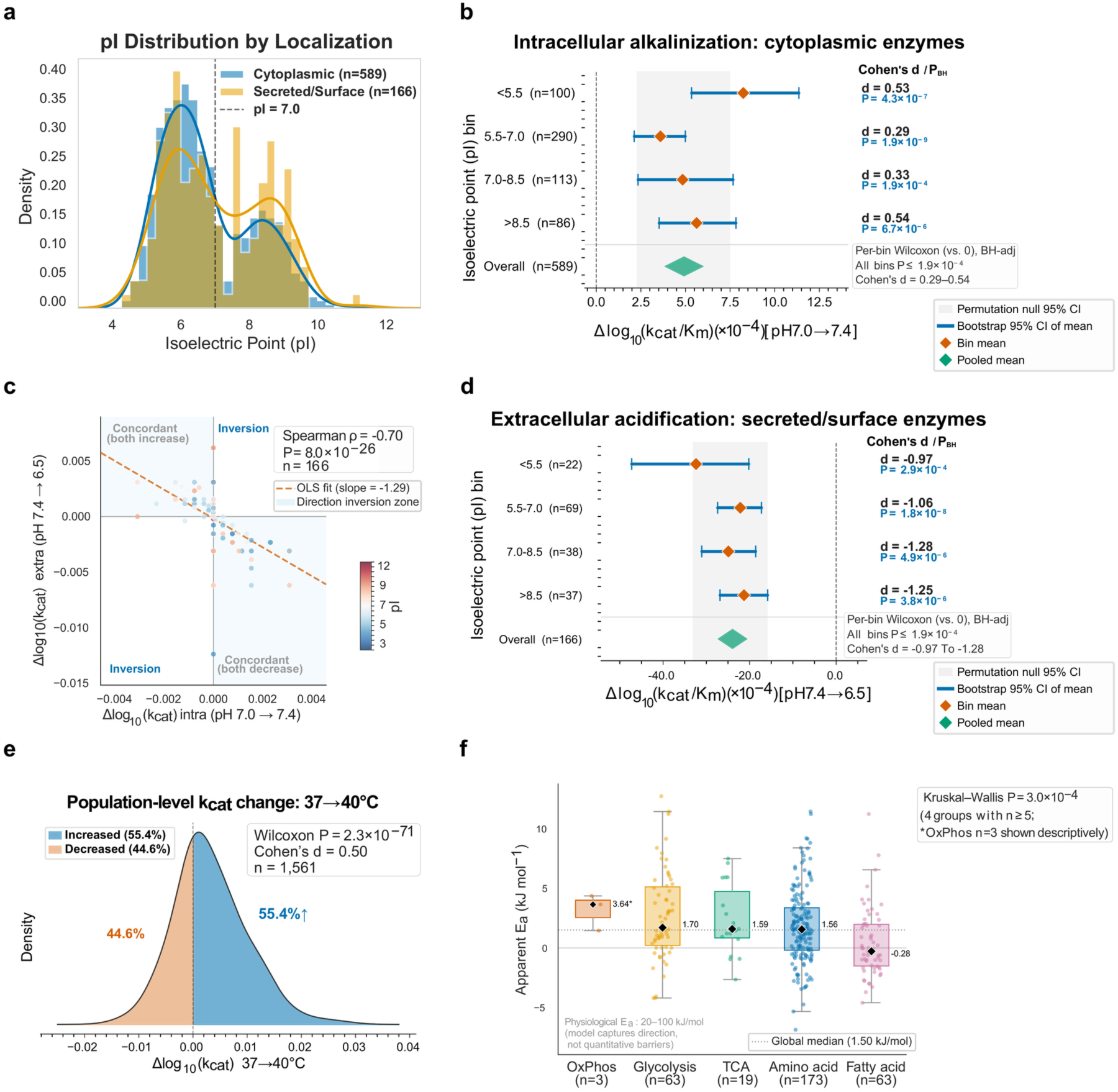
Condition-aware predictions capture biophysically consistent kinetic responses. a, Distribution of theoretical isoelectric points for cytoplasmic (*n* = 589) and secreted/extracellular (*n* = 166) human enzymes. b, Change in predicted log_10_(*k*_cat_/*K*_*m*_) under intracellular alkalinization (pH 7.0 → 7.4) stratified by pI bin; BH-adjusted *P* ≤ 1.9 × 10^−4^ across bins. c, Direction inversion: intracellular vs. extracellular *k*_cat_ shifts for 166 secreted/extracellular enzymes (Spearman *ρ* = −0.70, *P* = 8.0 × 10^−26^). d, Change in predicted log_10_(*k*_cat_/*K*_*m*_) under extracellular acidification (pH 7.4 → 6.5), showing the opposite direction across bins. e, Distribution of *Δ*log_10_(*k*_cat_) from 37 to 40 °C across 1,561 human enzymes; 55.4% show increased *k*_cat_(one-sided Wilcoxon *P* = 2.3 × 10^−71^, Cohen’s *d* = 0.50). f, Distribution of model-derived apparent activation energies across metabolic pathways (Kruskal–Wallis *P* = 3.0 × 10^−4^); the panel notes the physiological range (20–100 kJ mol^−1^) for comparison.

For temperature, we scanned seven points (30–45°C) at pH 7.0. A majority of human enzymes (55.4%) showed increased *k*_cat_from 37 to 40°C (one-sided Wilcoxon *P* = 2.3 × 10^−71^, *d* = 0.50; Fig. 4e). Apparent activation energies (*E*_*a*_) estimated from Arrhenius regression differed across metabolic pathways (Kruskal–Wallis *P* = 3.0 × 10^−4^), with median *E*_*a*_ highest for oxidative phosphorylation (3.64 kJ mol^−1^), followed by glycolysis (1.70) and the TCA cycle (1.59) (Fig. 4f). The absolute *E*_*a*_magnitudes were attenuated relative to physiological values (20–100 kJ mol^−1^), indicating that EnzCast captures qualitative temperature direction and pathway stratification rather than quantitative activation barriers (Extended Data Fig. 13).

Together, these scans confirm that the condition encoder learned compartment-appropriate pH directionality (*ρ* = −0.70) and non-random temperature responses with detectable pathway stratification. These condition-aware priors provide the biochemical layer for the tumor-biology analyses below.

### 2.4 Patient-specific kinetic landscapes reveal cell-type-dependent metabolic rewiring

EnzCast provides reaction-level kinetic estimates but does not capture patient-specific variation arising from tumor-associated isoform usage. We therefore developed IsoKin, an isoform-resolved Bayesian calibration framework that combines EnzCast priors—generated from models retrained on the full KinBench dataset under physiological intracellular conditions (pH 7.0, 37°C)—with single-cell RNA-seq (scRNA-seq) data and Recon3D [20] stoichiometric constraints to infer apparent *in vivo* kinetic parameters at individual patient and cell-type resolution (Fig. 5). IsoKin expands the gene–protein–reaction (GPR) axis to include splicing isoforms and derives context specificity through expression-weighted effective kinetics, 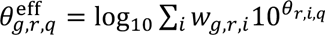 (METHODS).

**Fig. 5.**
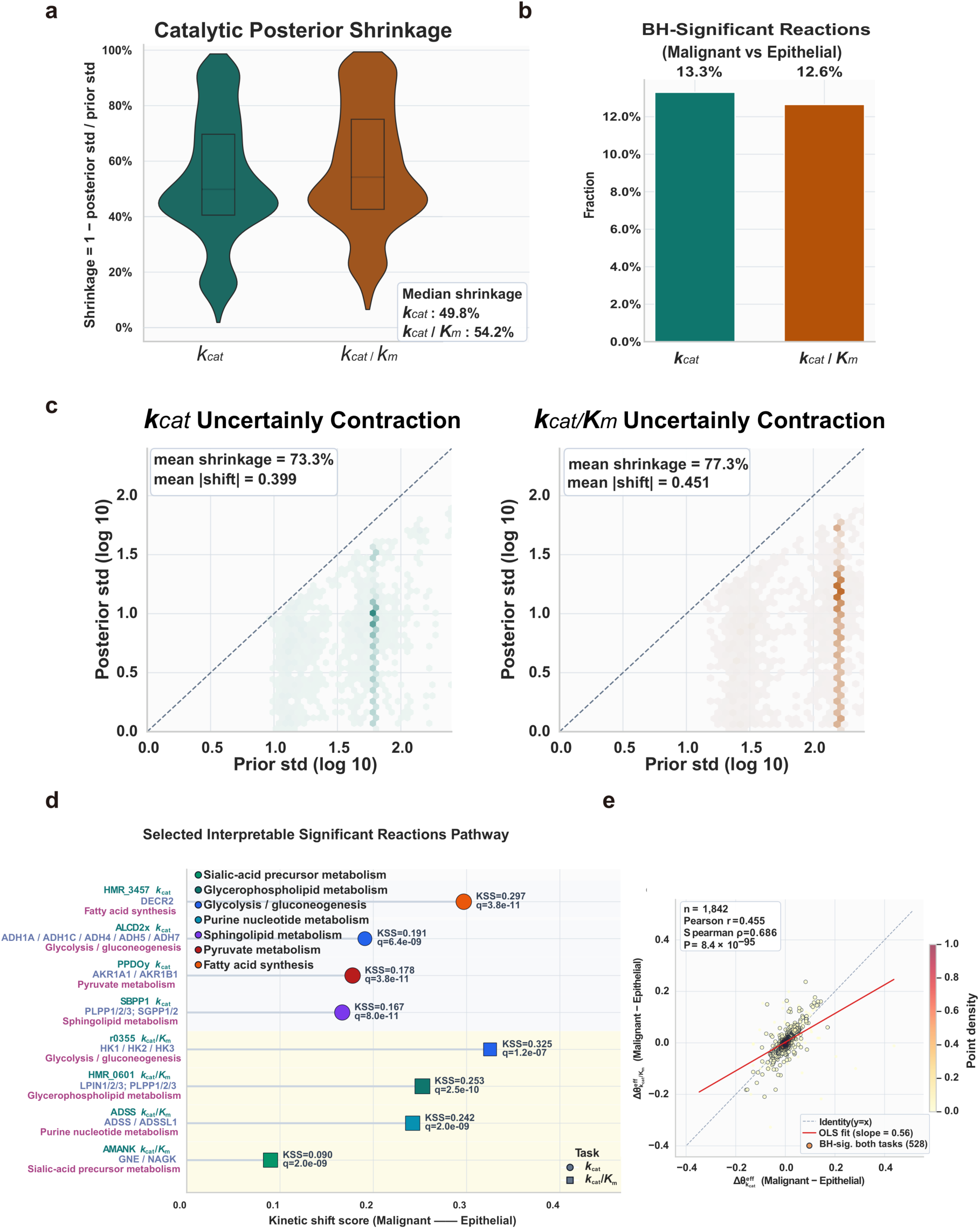
IsoKin calibrates EnzCast priors and highlights interpretable catalytic rewiring in the CRC cohort [25]. a, Posterior shrinkage (1 − *σ*_post_/*σ*_prior_) for *k*_cat_ and *k*_cat_/*K*_*m*_across 1,842 reactions; medians 49.8% and 54.2%. b, BH-significant ( FDR < 0.05) malignant-vs.-epithelial differences: 245 (13.3%) and 233 (12.6%). c, Prior versus posterior standard deviation; shrinkage 73.3% (*k*_cat_) and 77.3% (*k*_cat_/*K*_*m*_); shifts 0.399 and 0.451 log_10_units. d, Selected interpretable BH-significant reactions; point position shows kinetic shift score, color encodes metabolic subsystem, and q values are annotated. e, Cross-task *Δθ*^eff^ coherence (*r* = 0.455, *P* = 8.4 × 10^−95^; *ρ* = 0.686). Priors: pH 7.0, 37 °C.

Applied to a colorectal cancer (CRC) cohort of 20 patients and 422,861 cells [25], IsoKin resolved 1,842 metabolic reactions across 318 patient-by-cell-type contexts, of which 1,045 reactions (56.7%) contained multiple splicing isoforms. Posterior uncertainty decreased sharply for the catalytic parameters: mean posterior standard deviation fell by 73.3% for *k*_cat_and 77.3% for *k*_cat_/*K*_m_(Fig. 5a,c), with median reaction-level reductions of 49.8% and 54.2%, respectively. *K*_m_refinement was modest (16.4%). We therefore focused downstream analyses on *k*_cat_ and *k*_cat_/*K*_m_as the parameters most directly linked to catalytic throughput.

Malignant-vs.-epithelial comparisons identified BH-significant differences for 245 of 1,842 reactions (13.3%) for *k*_cat_ and 233 (12.6%) for *k*_cat_/*K*_m_(Fig. 5b). Reaction-level shifts in the two catalytic parameters positively correlated (Pearson *r* = 0.455, *P* = 8.4 × 10^−95^; Spearman *ρ* = 0.686; Fig. 5e), consistent with coherent but nonidentical metabolic rewiring. Among the BH-significant reactions, selected examples mapped to interpretable metabolic modules, including pyruvate metabolism (AKR1A1/AKR1B1), fatty-acid synthesis (DECR2), sphingolipid and glycerophospholipid remodeling (PLPP/SGPP and LPIN/PLPP), purine nucleotide metabolism (ADSS/ADSSL1), and sialic-acid precursor metabolism (GNE/NAGK) (Fig. 5d).

Notably, many differentially regulated reactions are net producers or consumers of protons, connecting to the pH-dependent responses in Section 2.3. The clustering of BH-significant kinetic shifts around H^+^-coupled reactions suggests that patient-specific isoform usage amplifies a coherent metabolic signature of intracellular pH adaptation.

The BH-significant reactions also included sialic acid metabolism at both pathway ends: biosynthesis (GNE/NAGK) and degradation by neuraminidases (NEU1–4). Because hyper-sialylation promotes immune evasion and metastasis, we tested whether neuraminidase kinetic shifts correlated with the metastatic phenotype. Across the 20-patient cohort, signed malignant-vs.-epithelial neuraminidase *k*_cat_ shifts correlated with epithelial–mesenchymal transition (EMT) scores (Spearman *ρ* = −0.54, BH *P* = 0.027; Extended Data Fig. 16a), with partial correlation confirming that this association is independent of oxidative stress ( *r* = −0.53, *P* = 0.020 after controlling for ROS; *r* = −0.003, *P* = 0.989 for ROS after controlling for EMT; Extended Data Fig. 16b). The negative sign indicates that high-EMT tumors exhibit more negative neuraminidase shifts, consistent with a reduced-degradation hypothesis for hyper-sialylation rather than biosynthesis upregulation.

Across the 318 patient-cell-type contexts (Fig. 6a), malignant cells diverged more substantially from stromal cells than from originating epithelial cells: BH-significant differential reactions for *k*_cat_ accounted for 38.3% in malignant-vs.-macrophage and 38.9% in malignant-vs.-fibroblast comparisons, but only 13.3% in malignant-vs.-epithelial comparisons (Extended Data Table 4). This approximately threefold larger differential against stromal cell types supports use of the originating epithelial lineage as the most informative control.

**Fig. 6.**
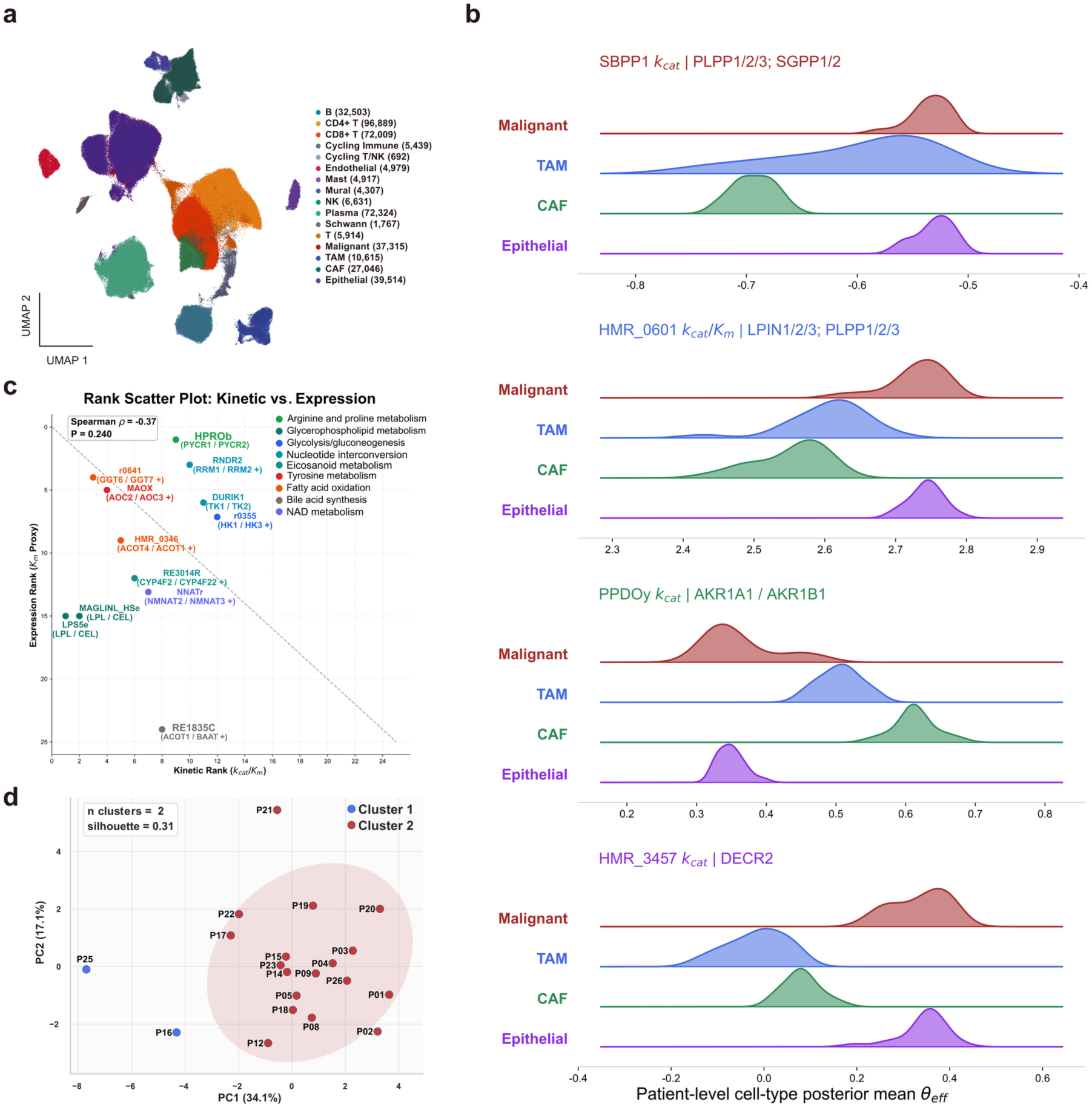
Cell-type kinetics and patient-specific rewiring reveal interpretable metabolic heterogeneity in the CRC cohort [25]. a, UMAP of 422,861 CRC cells from 20 patients (16 annotated types); Malignant, TAM, CAF, and Epithelial groups highlighted as primary IsoKin analysis groups. b, Patient-level posterior means per cell type for four BH-significant reactions (reaction ID; gene family): sphingolipid (SBPP1; PLPP/SGPP), glycerophospholipid (HMR_0601; LPIN/PLPP), pyruvate (PPDOy; AKR1A1/AKR1B1), and fatty-acid (HMR_3457; DECR1/DECR2) metabolism; HMR_3457, HMR_0601, and SBPP1 are elevated in malignant cells, PPDOy shows the opposite. c, Kinetic importance rank versus expression-only rank for the top-12 reactions; point color denotes metabolic subsystem, shape denotes task (circle: *k*_cat_; square: *k*_cat_/*K*_*m*_); Spearman *ρ* = −0.37, *P* = 0.240 (non-significant). d, PCA of the malignant-vs.-epithelial rewiring matrix; PC1 explains 34.1% of the variance, PC2 explains 17.1%, and the optimal two-cluster partition achieves a silhouette score of 0.31. SVI convergence diagnostics are shown in Extended Data Fig. 3b, and the full reaction ranking and patient-by-reaction heatmap are shown in Extended Data Fig. 4.

Within these BH-significant differential reactions, malignant-vs.-stromal kinetic shifts did not follow a single uniform direction. Three lipid- and sphingolipid-related reactions—the fatty-acid - *β* oxidation reaction HMR_3457 (DECR1/DECR2), the glycerophospholipid-remodeling reaction HMR_0601 (LPIN/PLPP), and the sphingolipid reaction SBPP1 (PLPP/SGPP)—were shifted upward in malignant cells relative to macrophages and fibroblasts, whereas the pyruvate-metabolism reaction PPDOy (AKR1A1/AKR1B1) showed the opposite direction (Fig. 6b). This reaction-specific reallocation of catalytic activity—toward lipid biosynthesis and away from pyruvate-aldol reduction—indicates that malignant kinetic remodeling is not uniform but module-specific. Patient-level heterogeneity in this rewiring was structured: PCA of the malignant-vs.-epithelial rewiring matrix separated patients into two broad groups (Fig. 6d).

To prioritize candidate vulnerabilities, we ranked reactions by a composite kinetic importance score that integrates malignant-vs.-epithelial shift magnitude, across-patient rewiring variability, and posterior information gain (METHODS). The top-ranked interpretable reactions spanned glycerophospholipid remodeling (LPS5e and MAGLINL_HSe), fatty-acid oxidation (r0641 and HMR_0346), tyrosine metabolism (MAOX, AOC1/AOC2/AOC3), eicosanoid metabolism (RE3014R), NAD metabolism (NNATr (NMNAT1)), bile acid synthesis (RE1835C), arginine and proline metabolism (HPROb (PYCR2)), and nucleotide interconversion (RNDR2 (RRM1/RRM2) and DURIK1 (TK1/TK2)) (Fig. 6c; Extended Data Fig. 11a). This ranking did not collapse onto an expression-only control ranking (Spearman *ρ* = −0.37, *P* = 0.240; Fig. 6c), indicating that isoform-specific kinetic differences and flux-balance constraints contribute information at least partly independent of transcript abundance.

Beyond the IsoKin posteriors, condition-aware scans identified two additional patterns: (i) xenobiotic-metabolism enzymes showed significantly greater catalytic efficiency downregulation under intracellular alkalinization than the metabolic background (BH *P* = 0.007; Extended Data Fig. 14), suggesting a kinetic complement to multidrug resistance; and (ii) energy-production pathway enzymes showed amplified temperature responses (Mann–Whitney *P* = 8.3 × 10^−6^; Extended Data Fig. 15), with scRNA-seq evidence of UCP4/UCP5 upregulation in malignant cells supporting a thermogenesis-enhanced-catalysis hypothesis (Supplementary Note 1).

Together, these results show that IsoKin-derived posterior kinetics, combined with condition-aware scans, expose a layer of tumor heterogeneity partly orthogonal to transcript abundance and generate interpretable reaction-level hypotheses.

### 2.5 Orthogonal validation provides mixed support for inferred kinetic rewiring

To assess whether IsoKin-inferred kinetic rewiring reflects genuine metabolic differences, we compared posterior kinetics against three independent platforms not used in constructing the posterior. First, we compared IsoKin effective *k*_cat_/*K*_m_values with metabolic flux scores from scFEA [26], an independent single-cell flux estimation framework applied to the same CRC cohort (METHODS).

Across the full mapping set (833 reactions, 33,320 matched context-reaction pairs across 20 patients), posterior *k*_cat_/*K*_m_ values correlated more strongly with scFEA flux scores than the matched EnzCast priors (posterior Pearson *r* = 0.050 vs. prior *r* = 0.033; Steiger *P* = 0.0046; Fig. 7a). A conservative high-confidence subset (single selected scFEA module per reaction; mean mapping F1 ≥ 0.5; 246 reactions, 9,840 matched context-reaction pairs) yielded a directionally consistent but non-significant posterior-over-prior advantage (posterior *r* = 0.074; prior *r* = 0.059; Steiger *P* = 0.141; Fig. 7b), whereas a within-patient malignant-vs.-epithelial delta analysis confirmed that cell-type-specific kinetic rewiring aligns with flux differences (*r* = 0.049, *P* = 0.0005; Fig. 7c). Together, the posterior-over-prior advantage reaches significance as mapping coverage increases.

**Fig. 7.**
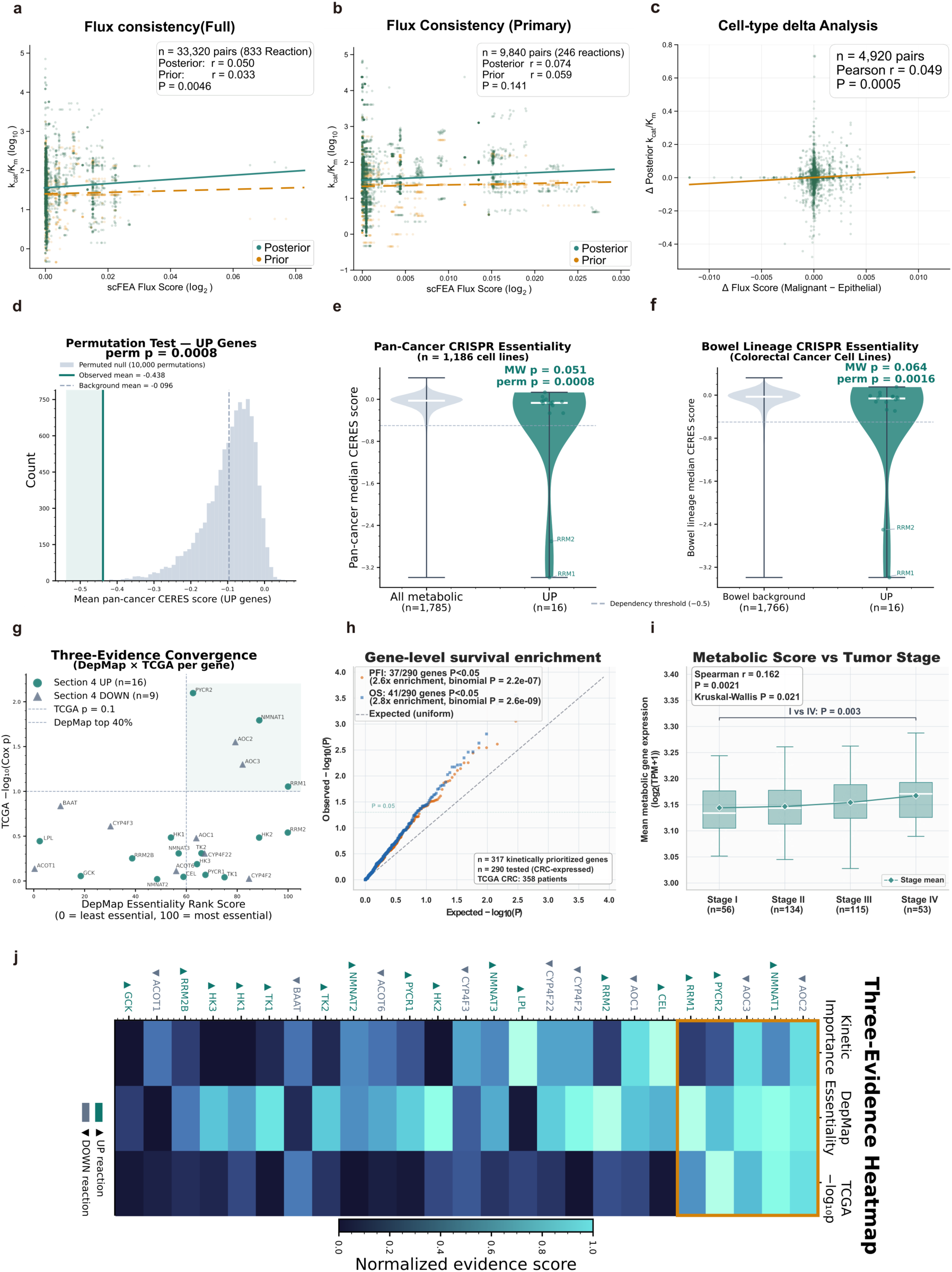
Orthogonal validation of inferred kinetic rewiring. a–c, scFEA flux-consistency: posterior versus prior *k*_cat_/*K*_*m*_ correlation for the full set (*r* = 0.050 vs *vs*0.033, Steiger *P* = 0.0046; a), primary subset (*r* = 0.074 vs *vs*0.059, *P* = 0.141; b), and within-patient delta (*r* = 0.049, *P* = 0.0005; c). d–f, DepMap essentiality: permutation null (d), pan-cancer (e), and bowel-lineage (f) CERES scores. g, Three-platform convergence scatter (5 genes: AOC2, NMNAT1, AOC3, PYCR2, RRM1). h,i, TCGA CRC survival enrichment QQ plot (h) and composite metabolic score versus AJCC tumor stage (Spearman *r* = 0.162, *P* = 0.0021; i). j, Three-evidence convergence heatmap.

We next tested the 16 genes catalysing the top-ranked kinetically up-regulated reactions from Section 2.4 against 1,186 DepMap [27] 24Q4 cancer cell lines spanning a metabolic background of 1,785 genes (METHODS). These genes showed substantially lower mean pan-cancer CERES scores than the metabolic background (mean = −0.438 vs. −0.096; one-sided Mann–Whitney *P* = 0.051; permutation *P* = 0.0008, 10,000 permutations; Fig. 7d,e), a signal driven substantially by RRM1 (CERES = −3.39) and RRM2 (CERES = −2.70), both well-established pan-cancer essential genes whose high kinetic importance as predicted by IsoKin is concordant with their observed pan-cancer dependency. The signal was preserved in the bowel cancer lineage (Mann–Whitney *P* = 0.064; permutation *P* = 0.0016; Fig. 7f).

To further strengthen the functional interpretation, we integrated three independent evidence streams—kinetic importance rank, per-gene CRC TCGA overall survival Cox *P*-value, and pan-cancer DepMap essentiality rank—into a convergence analysis. Five genes satisfied all three criteria (DepMap rank > 60th percentile and TCGA Cox *P* < 0.1): AOC2 (CERES = −0.143, Cox *P* = 0.028), NMNAT1 (CERES = −0.264, Cox *P* = 0.016), AOC3 (CERES = −0.169, Cox *P* = 0.050), PYCR2 (CERES = −0.065, Cox *P* = 0.008), and RRM1 (CERES = −3.391, Cox *P* = 0.088; Fig. 7g,j). Among these, PYCR2 (proline biosynthesis) ranks at the 63rd DepMap percentile with TCGA OS Cox *P* = 0.008, and NMNAT1 (NAD^+^biosynthesis) reaches the 89th percentile with Cox *P* = 0.016, providing convergent multi-platform support for their kinetic prioritization.

Finally, we tested whether the 317 kinetically prioritized genes are enriched for survival associations in an independent TCGA [28] CRC cohort (*n* = 377; progression-free interval, 103 events; overall survival, 85 events). After filtering to genes expressed in the TCGA CRC cohort, 290 of 317 candidates remained testable. Of these, 37 reached *P* < 0.05 for progression-free interval (expected 14.5 under the null; 2.6-fold enrichment; binomial *P* = 2.2 × 10^−7^) and 41 for overall survival (2.8-fold; *P* = 2.6 × 10^−9^; Fig. 7h); Kolmogorov–Smirnov tests confirmed departure from the uniform null (*P* < 10^−4^ for both endpoints). The directional split was balanced (18 risk vs. 19 protective for PFI; 20 vs. 21 for OS), consistent with metabolic rewiring involving both pro-tumorigenic and protective functions.

Independently, among the *n* = 358 patients with available American Joint Committee on Cancer (AJCC) staging, a composite metabolic score computed as the mean log_2_-transformed expression of the 290 kinetic genes increased monotonically with AJCC tumor stage (Spearman *r* = 0.162, *P* = 0.0021; Kruskal–Wallis *P* = 0.021; stage I vs. stage IV Mann–Whitney *P* = 0.003; Fig. 7i), indicating that aggregate metabolic gene expression among kinetically prioritized genes tracks with disease progression.

Taken together, the orthogonal evidence converges on a consistent directional signal despite mixed significance levels: posterior-over-prior scFEA advantage reaches significance in the full mapping set (Steiger *P* = 0.0046) but not the conservative subset (*P* = 0.141); DepMap essentiality is permutation-significant (*P* = 0.0008) but driven substantially by RRM1/RRM2. The convergence of five genes across three platforms and significant TCGA survival enrichment support the biological relevance of IsoKin-inferred kinetic rewiring.

## 3 DISCUSSION

EnzCast with IsoKin closes a persistent gap between population-level kinetic prediction and patient-specific apparent parameters. At the benchmark level, KinBench provides a unified four-task resource spanning 359,652 curated measurements, and EnzCast achieves task-adaptive prediction accuracy validated by systematic condition scans that confirm biophysically grounded pH and temperature responses (*ρ* = −0.70 direction inversion). At the patient-specific level, IsoKin converts those priors into effective reaction kinetics through isoform-resolved stoichiometric constraints and single-cell transcriptomic evidence, sharply reducing catalytic-parameter uncertainty (73.3% posterior shrinkage for *k*_cat_ and 77.3% for *k*_cat_/*K*_m_). Orthogonal analyses provided mixed but directionally consistent support: posterior kinetics correlated with independently estimated scFEA fluxes (Steiger *P* = 0.0046 in the full mapping set), kinetically up-regulated genes showed permutation-significant DepMap essentiality (*P* = 0.0008) with five genes converging across three platforms, and TCGA survival enrichment was significant for both endpoints, establishing EnzCast as a route from biochemical priors to patient-level metabolic hypotheses.

This positioning distinguishes EnzCast from earlier learning-based kinetic predictors, which are typically evaluated as stand-alone regressors without a mechanism for converting predictions into context-specific apparent in vivo parameters. In EnzCast, experimental conditions are introduced during prediction and uncertainty is carried forward into the Bayesian model. In IsoKin, expression-derived isoform fractions weight each isoform’s contribution to context-specific effective kinetics, with inference constrained by stoichiometric flux balance. The resulting representation provides a mechanistic layer between raw expression profiles and downstream metabolic interpretation.

The condition-aware architecture represents a qualitative advance over existing predictors, all of which treat experimental conditions as fixed or absent. The condition scans revealed biophysically grounded relationships learned from training data alone, including compartment-specific pH direction inversion and pathway-level temperature heterogeneity. Applied to tumor biology, these predictions identified kinetic mechanisms linking alkalinization to xenobiotic-metabolism downregulation, amplified energy-pathway catalysis under elevated temperature, and neuraminidase kinetic suppression in high-EMT tumors consistent with hyper-sialylation. However, attenuated temperature response magnitudes and the limited cohort size (*n* = 20) indicate that larger training datasets and expanded clinical cohorts will be needed to realize the full quantitative potential of this approach.

The colorectal cancer analyses reinforce this point. The approximately threefold larger kinetic differential against stromal cell types relative to originating epithelial cells argues against uniform malignant upregulation and instead points to reaction-specific reallocation of catalytic activity. The modest negative correlation between kinetic importance and expression-only rankings (*ρ* = −0.37, *P* = 0.240) is consistent with partial independence, suggesting that kinetic inference and transcript abundance capture at least partially non-redundant axes of metabolic variation.

Beyond the colorectal cancer application, EnzCast is available as a public webserver (https://enzcast.med.sustech.edu.cn) that accepts protein sequences, substrate SMILES, and optional experimental conditions, returning predicted *K*_m_, *k*_cat_, *k*_cat_/*K*_m_, and *K*_i_ values with uncertainty estimates. Batch submissions and CSV export are supported for downstream metabolic modelling.

Two principal limitations should be noted. First, the biological analyses are anchored to a single CRC cohort, demonstrating within-cohort heterogeneity rather than pan-cancer generality. Second, the orthogonal evidence is directionally consistent but not uniformly strong, with the conservative scFEA subset and DepMap signal each carrying specific caveats (Section 2.5). The current study therefore establishes a framework and mechanistic candidates rather than definitive clinical validation.

Future work should proceed along two coupled directions. On the modeling side, incorporating additional evidence layers such as proteomics, metabolomics, or direct flux readouts could further constrain patient-specific posteriors. On the translational side, the most pressing need is broader validation across independent cohorts, additional tumor types, and prospective tests of whether inferred kinetic rewiring predicts metabolic vulnerability or drug response. With directionally consistent validation outcomes across three independent platforms, the present framework offers a structured starting point for moving from sequence-scale enzyme knowledge to patient-specific metabolic interpretation.

## 4 METHODS

### 4.1 Data collection and KinBench benchmark construction

We constructed the KinBench benchmark by integrating measurement data of enzyme kinetics from four sources: SABIO-RK [22] and BRENDA [23], two manually curated repositories of biochemical reaction kinetics, supplemented with curated records from the TurNuP and CatPred published datasets generated using machine learning methods. The benchmark covers four kinetic parameters: Michaelis constant (*K*_m_, mM), turnover number (*k*_cat_, s^−1^), catalytic efficiency (*k*_cat_/*K*_m_, mM^−1^ s^−1^), and inhibition constant (*K*_i_, mM), comprising 181,663 *K*_m_, 99,173 *k*_cat_, 39,799 *k*_cat_/*K*_m_, and 39,017 *K*_i_ measurements (359,652 records in total) (Fig. 1, Module 1).

#### Substrate standardization

All substrate identifiers were converted to canonical SMILES IDs using RDKit [29]. During post-merge standardization, substrate names that appeared under multiple synonyms were unified by mapping each synonym to its most frequently observed SMILES ID in the merged dataset; when frequency was tied, the entry with the highest annotation completeness (valid SMILES, standard units, and UniProt-mapped enzyme) was selected as the representative structure. When a substrate name resolved to multiple candidate SMILES IDs—for example, the same trivial name maps to structurally distinct compounds across different source databases—cofactor-like candidates (ATP, NAD^+^, CoA, and their common derivatives) were excluded from the representative selection, so that the retained identifier corresponds to the biologically relevant small-molecule substrate rather than a ubiquitous cofactor. Each retained substrate annotation was then standardized to a single canonical SMILES ID using RDKit’s canonicalization algorithm, which normalizes atom ordering, stereochemistry conventions, and aromaticity perception, yielding a unique, graph-constructible structure for downstream molecular graph encoding.

#### Protein sequence processing

Protein identifiers were mapped to UniProtKB [24] accession IDs. For entries lacking a direct UniProt mapping, we performed cascading API queries using Enzyme Commission (EC) numbers and NCBI taxonomy identifiers. Sequences were normalized by converting to uppercase, removing space and stop codons. For proteins exceeding the 1,024-residue input length of the sequence encoder, we applied an active-site-centered cropping strategy with a three-tier fallback: the crop center was determined from (1) UniProt functional annotations (active site, binding site, metal-binding, nucleotide-binding, or generic site residues), (2) the structural centroid of the C_*α*_ atoms with predicted local distance difference test (pLDDT) scores ≥ 70, or (3) the sequence midpoint as the last resort. A symmetric window of 1,024 residues centered on this position was then extracted, preserving catalytically relevant spatial context.

#### Structural feature enrichment

Three-dimensional protein structures were obtained in a tiered manner: experimental structures from the Protein Data Bank (PDB) [30] have the highest priority for inclusion, followed by AlphaFold [31] predictions with pLDDT ≥ 70, and ESMFold [14] for de novo folding of remaining sequences. Protein structure graphs were constructed with C_*α*_ atoms as nodes (21 amino acid type embeddings) and *k*-nearest-neighbor edges (*k* = 10, a neighborhood size that captures both direct contacts and second-shell interactions within ∼ 10Å and is standard in protein structure graph neural networks) with Euclidean distances as the edge weights. Substrate molecular graphs were derived from SMILES, having atoms as nodes with 42-dimensional feature vectors (20-dimensional atom-type one-hot encoding, 5-dimensional chirality encoding, 8-dimensional degree one-hot encoding, 6-dimensional hybridization one-hot encoding, and 3 scalar features covering formal charge, aromaticity, and ring membership) and bonds as edges with 6-dimensional feature vectors.

#### Quality control

We removed entries with non-positive kinetic values, invalid molecular structures (failed RDKit parsing), or missing essential fields (UniProt identifier, amino acid sequence, or kinetic measurement). Records were deduplicated using a composite five-element key (EC number, organism, substrate name, measured value rounded to four significant figures, and UniProt identifier); when multiple records shared the same key, we retained the entry with the highest annotation completeness as assessed by a weighted quality score prioritizing valid UniProt mapping, sequence length, and SMILES completeness. All kinetic values were *log*_10_-transformed to compress the dynamic range spanning over 18 orders of magnitude. pH and temperature values were extracted from free-text annotation fields via regular expressions; ranges were converted to arithmetic means.

#### Benchmark partitioning

To ensure that benchmark evaluation reflects generalization to genuinely novel enzyme families rather than memorization of homologous training data, we adopted a sequence-similarity-based partitioning strategy. All unique protein sequences were clustered at 50% sequence identity using MMseqs2 easy-linclust [19] with 80% bidirectional coverage (cov-mode 0), yielding 13,529 non-redundant clusters. Clusters were then partitioned into training, validation, and test sets using group-aware splitting, ensuring very low sequence-homology overlap between partitions. This design guarantees that every test-set enzyme belongs to a protein family entirely absent from the training data, directly testing whether models can extrapolate to evolutionarily distant enzyme families—the generalization scenario most relevant to enzyme discovery and patient-specific applications.

### 4.2 EnzCast model overview

EnzCast takes four inputs for each enzyme–substrate example: the amino acid sequence, a protein structure graph, a substrate molecular graph, and the associated experimental information (Fig. 1, Module 2). These modalities were encoded independently, and then integrated through substrate-to-structure cross-attention, condition-dependent modality scaling, and a gated fusion block, and decoded by a dual-pathway head that separates binding-related outputs (*K*_m_ and *K*_i_) from catalytic outputs (*k*_cat_ and *k*_cat_/*K*_m_). Both the multi-task and single-task training regimes described below shared this network architecture. Each task head returned a mean *μ* and log-variance log*σ*^2^, yielding heteroscedastic uncertainty estimates that were carried forward as priors in downstream Bayesian inference.

### 4.3 Protein sequence encoding

We used ESM-2 (650M parameters) [14] as the protein language model (PLM) backbone. To adapt the pre-trained PLM to kinetic parameter prediction while preserving its learned representations, we applied Low-Rank Adaptation (LoRA) [15] to the query, key, and value projection matrices of each transformer layer, with rank *r* = 8, scaling factor *α* = 16, and dropout rate 0.1. All other ESM-2 parameters remained frozen during training.

Rather than relying solely on the final ESM-2 layer, we extracted the hidden states from layers 10, 21, and 32 of the 33-layer model. Each layer output was independently normalized with LayerNorm, and the three tensors were combined through a learnable softmax-weighted sum. This multi-layer design preserves information across shallow motif-sensitive representations, intermediate domain-level features, and the broader functional context encoded in deeper layers.

The combined ESM-2 representation (1,280 dimensions) was projected to the working dimension (256) with a projection MLP: 1,280 → 512 → LayerNorm → GELU → Dropout(0.15) → 256 → LayerNorm → GELU → 256. Attention pooling over residue positions produced a global sequence summary 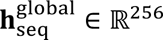. Input sequences were tokenized with the ESM-2 tokenizer and truncated or padded to a maximum length of 1,024 tokens.

### 4.4 Protein structure encoding

To capture the three-dimensional spatial relationships and geometric features not accessible from sequence alone, we designed a hierarchical virtual-node equivariant graph neural network (HVN-EGNN) [16].

The encoder comprised four stacked HierarchicalVNLayer modules [32], each combining two components. The first component, an equivariant graph convolutional layer (EGCL), updated both node features and three-dimensional coordinates. Edge messages were computed by an MLP operating on concatenated source and target node features, pairwise squared distance, and edge attributes; node features were updated by aggregating incoming edge messages weighted by learned attention coefficients. Coordinate updates were derived from interatomic displacement vectors scaled by feature-dependent weights, preserving equivariance to rigid-body rotations and translations. The second component, a virtual node layer, maintained a single learnable global node that aggregated information from all residues via multi-head attention (4 heads, head dimension 64) with a learnable temperature per head. The virtual node then propagated its updated representation back to all residues through a residual connection, and its coordinates were iteratively refined from attention-weighted residue displacements around the current graph centroid.

Node features were initialized from amino acid type embeddings (21 types → 256 dimensions), and edge features from edge-type embeddings, where the three types correspond to sequential backbone bonds, radius-based spatial contacts, and k-nearest-neighbor edges (each embedded into 256 dimensions). A sigmoid gate modulated the final virtual-node projection before output.

The encoder produced two representations: per-residue structural features **h**_struct_ ∈ ℝ^**N**×256^ and a virtual-node graph-level representation 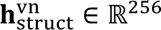, where **N** is the number of residues.

### 4.5 Substrate molecular encoding

Substrate molecular graphs were encoded in three stages. First, 42-dimensional atom features were projected to the working dimension (256) through a linear layer followed by LayerNorm and GELU activation. Second, three Graph Isomorphism Network (GIN) [17] layers with learnable **ε** parameters and residual connections refined the atom-level representations; each GIN layer used an internal MLP (256 → 512 → LayerNorm → GELU → 256). Third, a multi-head self-attention layer (4 heads) captured longer-range intramolecular dependencies across the propagated atom features.

To produce a graph-level molecular representation, we applied a multi-pooling strategy that combined four complementary pooling operations: global mean pooling (overall chemical context), global max pooling (salient functional groups), first-token embedding (anchor representation), and attention-weighted pooling (learned importance weighting). The four pooled representations were concatenated (4 × 256 = 1,024 dimensions) and projected to the working dimension via an MLP (1,024 → 512 → LayerNorm → GELU → Dropout(0.15) → 256).

In parallel, a 2,048-bit Morgan fingerprint [18] (radius 3) was computed from each substrate SMILES and projected to 128 dimensions through a separate MLP, providing a global chemical descriptor to the condition encoder.

The substrate encoder returned both atom-level features for downstream cross-attention and a graph-level embedding **h**_sub_ ∈ ℝ^256^.

### 4.6 Reaction condition encoding

Enzyme kinetic parameters are sensitive to experimental conditions, and the condition encoder integrates six sources of contextual information, each projected to a 128-dimensional embedding:

1. **Source organism**: learnable embedding lookup for each unique species in the training set. Including enzymes from diverse organisms substantially enlarges the training set and exploits the conservation of active-site geometry and substrate-binding thermodynamics across species. The organism embedding allows the model to learn species-specific offsets while sharing transferable physicochemical representations; for the downstream tumor application, all predictions were conditioned on *Homo sapiens*.
2. **Enzyme type**: three categories (wild-type, mutant, unspecified), each with a learnable embedding.
3. **pH**: a continuous value projected via MLP (1 → 64 → 128). A binary mask indicated missing values, which were replaced by a learnable default embedding.
4. **Temperature**: encoded similarly to pH.
5. **EC number**: learnable embedding initialized with Recon3D metabolic reaction embeddings when available, encoding prior knowledge about metabolic reaction similarity. Because Recon3D is a genome-scale reconstruction of *Homo sapiens* metabolism, these informed initializations covered 533 of the 3,972 unique EC numbers in the training set (13.4%), predominantly those catalyzing reactions in human central and intermediary metabolisms. This choice provides a biologically grounded inductive bias for the downstream application to human tumor metabolism, where the corresponding enzyme families are most clinically relevant. For the remaining EC numbers, embeddings were initialized from *N*(0, 0.02^2^), a small-variance distribution that keeps the initial representations near the origin and enables the model to learn task-relevant structure from data.
6. **Morgan fingerprint**: the 128-dimensional projection from the substrate encoder (Section 4.5). The six embeddings were concatenated (768 dimensions) and passed through a fusion MLP (768 → 256 → LayerNorm → GELU → Dropout → 128), yielding a condition vector **z**_cond_ ∈ ℝ^128^. A separate branch generated three adaptive scaling factors from **z**_cond_( 128 → 64 → SiLU → 3 → *σ*), where the terminal sigmoid constrains each factor to (0,1). These three scalar factors were broadcast across the sequence, structure, and substrate summary vectors, allowing experimental context to reweight the contribution of each modality. By setting pH, temperature, and organism to tumor-relevant values, the same encoder could be used to generate priors under cancer-specific conditions.

### 4.7 Multi-modal fusion and prediction heads

#### Substrate-to-structure cross-attention

To summarize enzyme–substrate interactions at residue resolution, a global substrate query was obtained by mean pooling the atom-level substrate features, and residue-level structural features served as keys and values in a 4-head cross-attention module. This operation produced a context-aware interaction summary **h**_cross_ ∈ ℝ^256^.

#### Adaptive feature scaling

The three sigmoid scaling factors from the condition encoder (Section 4.6) were broadcast-multiplied with the global sequence representation **h**^global^, the virtual-node structure representation 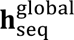, and the substrate representation **h**_sub_, respectively.

#### Gated fusion

The four scaled representations (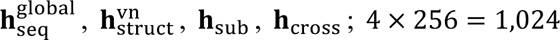 dimensions) were concatenated and processed by two parallel branches: a deep MLP (1,024 → 512 → LayerNorm → GELU → Dropout → 256 → LayerNorm → GELU) and a sigmoid gate (1,024 → 256 → *σ*). The MLP output was element-wise multiplied by the gate, producing a fused representation **h**_fused_ ∈ _ℝ_256.

#### Dual-pathway prediction head

The fused representation was concatenated with the condition vector ( 256 + 128 = 384 dimensions) and passed through a shared trunk ( 384 → 256 → LayerNorm → GELU → Dropout). Two specialized branches then produced predictions for kinetically related parameter groups:

- **Binding pathway** (*K*_m_ and *K*_i_): an MLP (256 → 128) followed by separate linear heads, each outputting a mean *μ* and log-variance log*σ*^2^.
- **Catalytic pathway** (*k*_cat_ and *k*_cat_/*K*_m_): a deeper MLP (256 → 256 → 128) followed by separate linear heads with the same output format.

This dual-pathway design reflects the biochemical distinction between binding equilibria (*K*_m_and *K*_i_) and catalytic throughput (*k*_cat_ and *k*_cat_/*K*_m_), encouraging shared internal representations within each pathway. The four output heads were present in both training regimes; regime-specific differences were introduced through the supervised objective and the choice of checkpoint family retained for each task.

### 4.8 Loss functions

**Beta-NLL loss:** The primary loss function was a variance-stabilized negative log-likelihood (Beta-NLL) [33]:

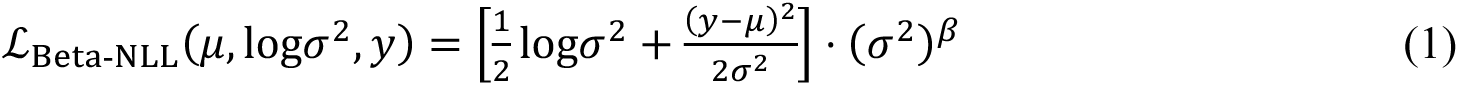

where *β* = 0.5 and the (*σ*^2^)^*β*^ weighting factor was applied with a stopped gradient to prevent the model from artificially reducing loss by inflating predicted variance. The log-variance was clamped to [−8,8] for numerical stability. To stabilize early training, the NLL component was weighted by a schedule that ramped linearly from 0.05 to 1.0 over the first 80% of training epochs, prioritizing mean prediction accuracy before variance refinement.

#### Uncertainty-weighted multi-task loss

In multi-task training mode, task losses were integrated using learned uncertainty weights following Kendall et al. [34]:

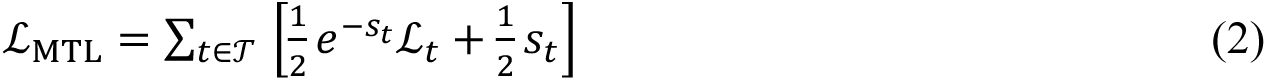

where *s*_*t*_ = log*σ*^2^_t_ are learnable log-variance parameters (clamped to [−6,6]) that automatically balance task difficulties, and *T* = {*K*_m_, *k*_cat_, *k*_cat_/*K*_m_, *K*_i_}.

**Consistency loss**: In multi-task mode, a soft constraint enforced the biochemical identity log_10_(*k*_cat_/*K*_m_) = log_10_(*k*_cat_) − log_10_(*K*_m_):

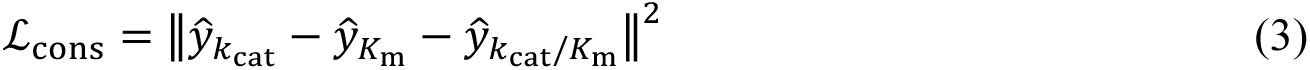

computed on denormalized log_10_-scale predictions, with weight *λ*_cons_ = 0.1.

### 4.9 Training procedure

We trained two model families (hereafter termed checkpoint families, referring to the distinct sets of saved model weights produced by each training regime) with identical encoders and the prediction-head topology. The multi-task family instantiated the full four-head architecture and used shared supervision during the supervised stage, whereas the single-task family optimized only one target head at a time. In the task-adaptive workflow used throughout the study, the catalytic outputs (*k*_cat_ and *k*_cat_/*K*_m_) were taken from the multi-task family, whereas *K*_m_ and *K*_i_were taken from dedicated single-task checkpoints. Both families used early stopping on validation *R*^2^, so the final checkpoints reflect the epoch at which generalization performance plateaued.

#### Multi-task training schedule

The multi-task training schedule comprised two stages:

*Stage 0: Contrastive pre-training* (3 epochs). A task-agnostic protein–substrate alignment objective trained the encoders to project matched enzyme–substrate pairs into a shared 128-dimensional space using NT-Xent loss [35] (symmetric cross-entropy with label smoothing 0.1). Learning rate: 5 × 10^−5^. All samples from all four tasks contributed regardless of the downstream prediction target.

*Stage 1: Multi-task supervised training* (5 epochs). The multi-task scripts instantiated four output heads and computed task-specific Beta-NLL losses on the corresponding samples, aggregated with the uncertainty-weighted objective ℒ_MTL_ and the consistency penalty *λ*_cons_ℒ_cons_. Differentiated learning rates were applied: 8 × 10^−5^ for the backbone and feature-extraction modules, and 6 × 10^−4^ for the condition encoder, fusion block and prediction heads. A cosine schedule [36] with 6% linear warmup governed the learning rate decay. Gradient accumulation over the 2 steps yielded an effective batch size of 44 (physical batch size 22 on a single NVIDIA A800 80 GB GPU). Gradients were clipped at a maximum norm of 1.0. Early stopping with patience of 2 epochs on validation mean *R*^2^ terminated training if no improvement was observed.

#### Single-task training

Single-task checkpoints shared the same encoder architecture and Stage 0 contrastive pre-training as the multi-task family. In the subsequent supervised stage, only the target head was active in the loss; the multi-task aggregation, uncertainty weighting, and consistency regularization were disabled, and the dataloader was restricted to samples of the target task. The contrastive pre-training stage retained the full unfiltered training set to preserve the encoder generality. All remaining hyperparameters—learning rates, gradient accumulation, and early-stopping criteria—were identical to the multi-task configuration.

#### Exponential moving average

An exponential moving average (EMA) of model weights (decay = 0.999) was maintained throughout training. Regular and EMA checkpoints were tracked in parallel during supervised training. For downstream prior generation, we used the Stage 1 EMA checkpoints from the full-dataset retraining scripts.

### 4.10 Isoform-resolved Bayesian posterior inference framework

We implemented IsoKin, a single-stage Bayesian flux model that combines EnzCast priors, gene–protein–reaction (GPR) logic, and Recon3D [20] stoichiometric constraints to calibrate isoform-level kinetic parameters (Fig. 1, Module 3). The inferred quantities represent apparent *in vivo* kinetics, reflecting effective catalytic behavior in the cellular microenvironment rather than purified enzyme constants. IsoKin operates on patient-by-cell-type groups derived from scRNA-seq and outputs posterior distributions for isoform-level kinetic parameters, from which context-specific effective kinetics are derived after inference. ***Isoform decomposition and group-level expression***: The Recon3D GPR rules were parsed and expanded to disjunctive normal form, such that each OR branch is for an isoform and each AND clause represents a complex of multiple protein units. For each patient-by-cell-type group **g**, scRNA-seq expressions were converted from log1p space back to a linear space, averaged within the group, and evaluated on each isoform branch. Single-gene branches used the corresponding group-mean expression, whereas multi-subunit branches were integrated by the GPR evaluator’s soft-min operation to approximate the rate-limiting subunit. Group-level isoform detection rates were defined as the fraction of cells in which a branch was active under the corresponding Boolean detection rule.

#### Isoform-level EnzCast priors

EnzCast priors were assigned at the isoform level by matching each reaction–isoform branch to precomputed EnzCast inference tables generated from full-dataset retraining under intracellular default conditions (pH 7.0, 37 °C, *Homo sapiens*, wild-type). Consistent with the task-adaptive benchmark selection described above, downstream prior generation used single-task checkpoints for *K*_m_ and *K*_i_, and multi-task checkpoints for *k*_cat_and *k*_cat_/*K*_m_. Matching proceeded hierarchically: an exact match on both reaction identifier and catalytic gene name was sought first; if that failed, a reaction-level average across all available isoform branches was used; and if both failed, a global mean prior derived from all reactions in the dataset served as the final fallback. For any non-exact assignment, the prior standard deviation was inflated by a factor of 1.5 to reflect the additional uncertainty introduced by inexact prior assignment. The constrained reaction set therefore included reactions supported by exact, reaction-average or global-fallback prior assignments after intersection with Recon3D coverage and isoform-expression support. All priors were represented in log_10_ space.

#### Flux-constrained IsoKin model

For each reaction *r*, isoform *i*, and kinetic task **q** ∈ {*K*_m_, *k*_cat_, *k*_cat_/ *K*_m_, *K*_i_}, isoform-level parameters were sampled as 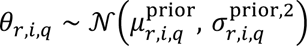. Reaction throughput depended on a reaction-level saturation index **ω**_*r*_ ∼ Beta(2,2)—a symmetric bell-shaped prior with mean 0.5 that encodes the expectation that intracellular reactions operate at intermediate substrate saturation rather than at boundary extremes—a substrate offset *s*_*r*_, and context-specific noise **ε**_g,*r*_, yielding

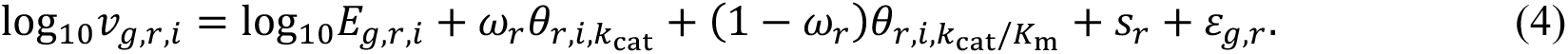

Reaction fluxes preserved OR semantics by summing isoform fluxes in linear space,

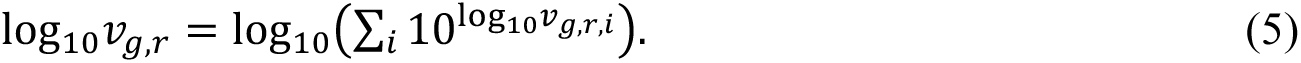

Low-detection contexts were down-weighted using a sigmoid reliability function of reaction detection rate, which reduced unreliable fluxes toward the reaction-wise median before evaluating the soft flux-balance likelihood. Net flux imbalance *Sv_g_* was penalized with a Gaussian tolerance term scaled by metabolite connectivity and the median estimated flux magnitude across all patient-by-cell-type groups for that metabolite. *K*_m_ was constrained indirectly through a soft consistency penalty enforcing *θ*_*r*,*i*,*k*cat_ − *θ*_*r*,*i*,*K*m_ ≈ *θ*_*r*,*i*,*k*cat/*K*m_, and *K*_i_ was sampled with a fixed, tight prior standard deviation of 0.2 (in log10 space) without a direct flux-balance signal. Because *k*_cat_ and *k*_cat_/*K*_m_are both directly constrained by the flux likelihood and are the parameters that most closely linked to catalytic throughput and metabolic reaction reprogramming, all downstream biological analyses in this work focus on these two posteriors as the primary IsoKin outputs.

Context-specific effective kinetics were derived from posterior samples as

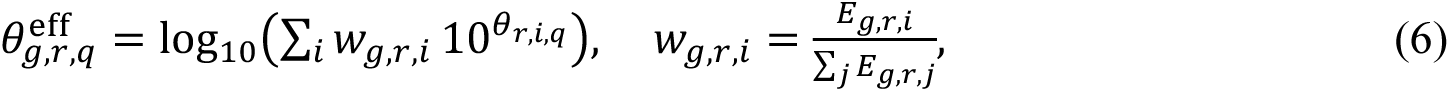

so that personalization emerged from context-specific isoform fractions rather than from an explicit reaction-level latent variable.

#### Inference and downstream metrics

Posterior inference used Pyro [37] with an AutoLowRankMultivariateNormal guide (rank = 10), ClippedAdam optimization, gradient clipping at 10, and a maximum of 20,000 SVI iterations. Early stopping was triggered when the relative loss over a 1,000-iteration window fell below 1 × 10^−3^. For the final CRC analyses, we used a learning rate of 0.001 and drew 2,000 posterior samples from the variational distribution to summarize the posterior means and standard deviations. IsoKin outputs were post-processed to compute the effective kinetics and variance decomposition across reactions and tasks. For the CRC comparisons reported in the main text, isoform switching scores (ISS) and kinetic shift scores (KSS) were derived from malignant-versus-epithelial contrasts in post-hoc analysis, with cell-type-specific differences assessed by two-sided Mann–Whitney *U* tests and Benjamini–Hochberg correction within each task and comparison.

For the analyses in Section 2.4, patient-specific rewiring was summarized as 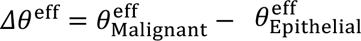 on the catalytic tasks. The detected rewiring converged on several metabolic axes linked to established cancer hallmarks [38]: glycerophospholipid remodeling and fatty-acid oxidation, which supply membrane building blocks and redox cofactors for rapid proliferation; nucleotide interconversion and NAD biosynthesis, which sustain the nucleotide and electron-carrier pools required by dividing cells; and eicosanoid metabolism, which modulates the inflammatory tumor microenvironment. Reactions were ranked by a composite kinetic importance score that combined normalized KSS, the across-patient standard deviation of *Δθ*^eff^, and mean posterior shrinkage weights (0.4, 0.3, and 0.3, respectively). Cell-type-specific differences were assessed with two-sided Mann–Whitney *U* tests followed by Benjamini–Hochberg correction within each task and cell-type comparison.

### 4.11 Condition scan protocol and database construction

#### Condition scan protocol

To evaluate the biophysical consistency of EnzCast’s condition encoder, we performed systematic pH and temperature scans across all 1,561 human enzymes in KinBench, using all three model configurations (multitask, single-task *K*_m_, single-task *K*_i_). For pH, seven conditions (6.5, 6.8, 7.0, 7.2, 7.4, 7.6, 7.8) were evaluated at 37°C. For temperature, seven conditions (30, 35, 37, 38.5, 40, 42, 45°C) were evaluated at pH 7.0. Per-enzyme kinetic shifts were computed as paired differences in log_10_space relative to the reference condition (pH 7.0 for pH scans; 37°C for temperature scans), then summarized as per-enzyme medians across substrate partners.

#### Database construction

Theoretical isoelectric points were computed using Biopython’s ProteinAnalysis.isoelectric_point() from full UniProt sequences (validated against ExPASy for 50 enzymes; Pearson *r* = 1.000). Subcellular localization was retrieved from the UniProt REST API (cc_subcellular_location field). Xenobiotic-metabolism enzymes (44 total: 24 Phase I, 20 Phase II) were curated from KEGG pathways hsa00980/hsa00982/hsa00983 and Recon3D. ATP-linked reactions (928 total) were identified from Recon3D reaction formulas containing ATP as substrate or product. Sialic acid pathway enzymes (29 total) were manually curated from KEGG and literature. Patient-level ROS and EMT scores were computed using decoupler’s run_ulm() on MSigDB HALLMARK gene sets from the CRC scRNA-seq dataset [25].

### 4.12 Single-cell RNA-seq data processing

IsoKin was applied to the CRC dataset [25] (422,861 cells from 20 patients and 16 annotated cell types, stored as an AnnData.h5ad object). Raw counts were processed with Scanpy [39] (v1.12). Quality control retained cells with 200–8,000 detected genes and 500–100,000 UMIs, excluded cells with more than 20% mitochondrial UMI fraction, as well as genes detected in fewer than three individual cells. Potential doublets were identified and removed using Scrublet [40] (v0.2.3). Batch effects across samples were corrected using Harmony [41], and unsupervised Leiden [42] clustering, followed by canonical-marker-based annotation of cells to 16 major types. Patient identifiers and harmonized cell-type annotations were extracted from the observation metadata, and raw integer counts were recovered from the original count layer for re-normalization.

The dataset was normalized to counts per million (target sum = 10^6^) and log(1 + *x*)-transformed. Unused auxiliary layers and annotations were removed before exporting a processed AnnData file for pipeline input. The processed object contained 320 patient-by-cell-type groups. The final IsoKin analysis retained 318 analyzable contexts after intersection with the constrained reaction set; the two excluded groups had insufficient expressed-gene overlap with the constrained metabolic reactions to support flux-constrained posterior inference. In downstream posterior analyses, malignant-vs.-epithelial comparisons contrasted malignant cells against epithelial cells according to the harmonized cell-type annotation.

### 4.13 Orthogonal validation analyses

#### Metabolic flux consistency

To assess whether IsoKin posteriors align with an independent estimate of a metabolic activity, we compared them against single-cell metabolic fluxes predicted by scFEA [26] on the same CRC expression matrix. The stored log1p-normalized expression values were first converted back to linear space (expm1) and averaged within each patient-by-cell-type group. The resulting pseudobulk profiles were restricted to the 651 human scFEA module genes present in the cohort and passed to the human M168 scFEA model (120 training epochs, no imputation).

Because scFEA reports module-level flux scores rather than individual Recon3D reaction fluxes, we established a mapping between IsoKin reactions and scFEA modules through shared enzyme genes. For each candidate reaction–module pair, gene-overlap coverage was defined as the fraction of the reaction’s associated genes present in the scFEA module, and specificity as the fraction of the module’s associated genes shared with the reaction (analogous to recall and precision, respectively). An F1-like composite was computed as the harmonic mean of coverage and specificity. We retained pairs with coverage ≥ 0.5, specificity ≥ 0.1, and an F1-like composite within 95% of the highest-scoring module for that reaction, and computed reaction-level scFEA scores as a weighted average of the selected module scores with weights proportional to the F1-like composite. To suppress ambiguous many-to-one mappings, the primary subset used in the main-text analysis was further restricted to reactions matched to a single scFEA module with mean mapping F1 ≥ 0.5.

We compared posterior and prior *k*_cat_/*K*_m_values against the matched scFEA scores, as this readout showed the clearest posterior–flux correlation under the high-confidence mapping filter. Pearson and Spearman correlations were computed across all matched context–reaction pairs, and posterior-versus-prior improvement was tested with the Steiger test for dependent correlations sharing the same scFEA response variable. As a sensitivity analysis for cell-type rewiring, we also correlated malignant-minus-epithelial posterior deltas with the corresponding scFEA deltas within each patient.

#### Genetic dependency validation

We tested whether genes encoding enzymes whose posterior catalytic parameters (*k*_cat_ or *k*_cat_/*K*_m_) were significantly elevated in malignant relative to epithelial cells (hereafter, kinetically up-regulated; Section 2.4) show stronger CRISPR essentiality using DepMap [27] 24Q4 gene-effect data (Supplementary Table X). The analysis focused on the 16 genes catalysing the top-ranked kinetically up-regulated reactions from the final 10 BH-significant reactions with small gene families (family size ≤ 4; Section 2.4). These genes were compared against a background of 1,785 metabolic-network genes across all 1,186 cancer cell lines in the DepMap 24Q4 release using a one-sided Mann–Whitney *U* test and an empirical permutation test (10,000 permutations). A separate bowel-lineage analysis applied the same test restricted to colorectal cell lines.

To identify genes supported by multiple independent lines of evidence, we performed a convergence analysis integrating three orthogonal data streams: kinetic importance rank (from Section 2.4), per-gene CRC overall survival univariate Cox *P*-value (computed as described below), and pan-cancer DepMap essentiality percentile rank. A gene was classified as convergent if it satisfied DepMap rank > 60th percentile and TCGA OS Cox *P* < 0.1.

#### Clinical outcome and gene-level enrichment

To evaluate whether genes associated with BH-significant differential reactions identified by IsoKin (Section 2.4; hereafter, kinetically prioritized genes) carry prognostic information, we performed a gene-level survival enrichment analysis in TCGA [28] colorectal adenocarcinoma (colon plus rectum primary-disease labels; *n* = 377 with available survival metadata). From the 317 unique genes supporting the 446 BH-significant reactions identified in Section 2.4, we retained those present in the TCGA expression matrix and expressed in CRC (TPM > 1 in > 10% of samples), yielding 290 evaluable genes.

For each gene, a univariate Cox proportional-hazards model was fitted on up to *n* = 377 samples with available survival metadata (371 for PFI, 373 for OS, after excluding samples with missing endpoint-specific values), using log_2_(TPM + 1) as the predictor against two endpoints: progression-free interval (PFI; 103 events) and overall survival (OS; 85 events). We counted genes reaching nominal significance (*P* < 0.05) and tested whether this count exceeded the 5% null expectation with a one-sided binomial test. Departure of the observed *P*-value distribution from uniformity was additionally confirmed with a Kolmogorov–Smirnov test.

For the stage association analysis, a composite metabolic score was computed as the mean log_2_(TPM + 1) across all 290 kinetic genes per sample. Its association with AJCC tumor stage (I–IV) was assessed by Spearman rank correlation, a Kruskal–Wallis test, and a one-sided Mann–Whitney *U* test comparing stage I versus stage IV. Samples with missing or discrepant stage annotations were excluded (*n* = 358 evaluable).

#### Web server implementation

The EnzCast web server provides a fully containerized inference environment for multi-modal kinetic parameter prediction. The backend integrates ESM-2 (650M parameters) for protein sequence encoding and the EnzCast graph neural network for substrate molecular graph and protein structure processing, executed through a FastAPI-based scheduling layer on a dedicated NVIDIA RTX 4090 GPU using bfloat16 mixed-precision arithmetic. A Gradio-based frontend accepts individual enzyme–substrate pairs or batch CSV submissions comprising UniProt accessions or raw sequences together with substrate SMILES strings and optional experimental conditions. For proteins exceeding 1,024 residues, the server applies the active-site-centered truncation strategy described above, using precomputed active-site centre coordinates to centre the context window on the catalytic pocket. Inference is executed deterministically. The server is available at https://enzcast.med.sustech.edu.cn; the full server source code is included in the EnzCast GitHub repository (https://github.com/Mxc666/EnzCast).

### 4.14 Baseline methods

We compared EnzCast with four published methods for enzyme kinetic parameter prediction.

**DLKcat** [10] encodes protein sequences with a convolutional neural network and substrate molecular graphs with a graph neural network; it was originally designed for *k*_cat_ prediction. We retrained DLKcat on the KinBench training set and evaluated on all applicable tasks.

**UniKP** [11] uses the ProtT5-XL protein language model [43] combined with pre-trained molecular representations and an ExtraTreesRegressor ensemble. It covers *k*_cat_, *K*_m_, and *k*_cat_/*K*_m_but does not use protein structure. We retrained UniKP on KinBench.

**TurNuP** [12] employs protein sequence and substrate features with a gradient-boosted ensemble architecture designed for *k*_cat_prediction. We retrained TurNuP on the KinBench training set using its default gradient-boosted ensemble architecture with ESM-1b sequence embeddings and structural fingerprint features.

**CatPred** [13] combines protein sequence and substrate SMILES with uncertainty estimation and covers *k*_cat_, *K*_m_, and *K*_i_ but not *k*_cat_/*K*_m_. We retrained CatPred on the KinBench training set.

All baselines were evaluated with the same train/validation/test splits, preprocessing, and metrics as EnzCast to ensure fair comparison. For tasks not natively supported by a baseline, performance was reported as not applicable (N/A). To verify implementation fidelity, we retrained each baseline on its original benchmark dataset using our reimplementation and compared the resulting metrics with originally reported values (Extended Data Table 5). UniKP reproduced the reported *K*_m_ and *k*_cat_/*K*_m_ *R*^2^ values within 0.03 and 0.06, respectively, and its *k*_cat_ PCC (0.81) was close to the reported 0.85. DLKcat matched the reported PCC (0.71), TurNuP *k*_cat_ exceeded the originally reported *R*^2^(0.637 vs 0.44, Pearson *r* = 0.81), and CatPred, evaluated in simplified single-model mode, remained within 0.03–0.10 *R*^2^of the reported ensemble values. These results support implementation fidelity while preserving the documented caveats for DLKcat regularisation details, CatPred ensemble/MVE loss, and TurNuP embedding extraction.

Unless noted otherwise, hyperparameter settings for each baseline followed those reported in the respective original publications.

### 4.15 Evaluation metrics and statistical analysis

All prediction metrics were computed in the log_10_ space, matching the training target representation.

#### Prediction performance

For benchmark evaluation, we computed five metrics in log_10_ space: coefficient of determination (*R*^2^), root mean squared error (RMSE), mean absolute error (MAE), Pearson correlation, and Spearman rank correlation. All multi-seed experiments used three independent random seeds (42, 123, and 2025), and summary statistics are reported as mean ± s.d. across seeds.

#### Statistical tests

Exact *P* values are reported throughout with the name and sidedness of the statistical test. Group comparisons in the IsoKin analyses used two-sided Mann–Whitney *U* tests followed by Benjamini–Hochberg (BH) correction [44] where applicable. Flux-consistency analyses used Pearson and Spearman correlations together with the Steiger test for dependent-correlation comparisons sharing the same response variable. DepMap enrichment analyses used one-sided Mann–Whitney *U* tests (kinetically up-regulated genes more essential than metabolic background) and empirical permutation tests (10,000 permutations). TCGA gene-level analyses used univariate Cox proportional-hazards models, with enrichment over the null quantified by a one-sided binomial test; *P*-value uniformity was assessed by a Kolmogorov–Smirnov test; stage association was evaluated by Spearman rank correlation and Kruskal–Wallis tests; and pairwise stage comparisons used one-sided Mann–Whitney *U* tests.

## SUPPLEMENTARY INFORMATION

Supplementary Information is available for this paper.

## ACKNOWLEDGMENTS

We thank the maintainers of SABIO-RK, BRENDA, DepMap, and TCGA, and the authors of the published colorectal cancer single-cell transcriptomic datasets analyzed here, for making these resources publicly available. We also acknowledge the open-source software communities behind PyTorch, PyTorch Geometric, Hugging Face Transformers, RDKit, and MMseqs2, as well as the GPU computing infrastructure that supported model training and Bayesian inference.

## DECLARATIONS

- **Funding:** This work was supported by the National Natural Science Foundation of China (NSFC, No. T2350010) and the National Natural Science Fund for International Scientists (No. W2431059). We also acknowledge the support of the Key University Laboratory of Metabolism and Health of Guangdong, Southern University of Science and Technology (No. 2022KSYS007), and the Center for Computational Science and Engineering at Southern University of Science and Technology. Additionally, we express gratitude to the SUSTech Homeostatic Medicine Institute, School of Medicine, Southern University of Science and Technology, Shenzhen 518055, China, for their support.
- **Competing interests:** The authors declare no competing interests.
- **Ethics approval:** This study used only publicly available, de-identified data and involved no new experiments on humans or animals. Accordingly, institutional review board approval and informed consent were not required.
- **Data availability:** Raw kinetic measurements were obtained from SABIO-RK[22] and BRENDA[23]. External validation data were obtained from previously published colorectal cancer single-cell RNA-seq datasets, DepMap 24Q4, and the TCGA colon and rectum adenocarcinoma cohort; accession details are provided in the Methods. The KinBench benchmark splits (training, validation, and test CSV files) are deposited in the EnzCast public repository at https://github.com/Mxc666/EnzCast. Pre-trained EnzCast model checkpoints are hosted on the Hugging Face Hub at https://huggingface.co/Mxc666/EnzCast. IsoKin posterior outputs for the displayed CRC cohort and the source tables underlying the main and Extended Data figures are available from the corresponding author upon reasonable request.
- **Code availability:** All model training, inference, and figure-generation scripts used in this study are publicly available at https://github.com/Mxc666/EnzCast. The repository includes the code for reproducing benchmark training and evaluation, the IsoKin inference pipeline, and the EnzCast web server source code. The web server is publicly accessible at https://enzcast.med.sustech.edu.cn.
- **Author contributions:** X. Mu: Conceptualization, Formal analysis, Methodology, Software, Investigation, Visualization, Writing – original draft, Writing – review & editing. Y. Yang: Visualization, Data curation, Writing – review & editing. Q. Wang: Visualization, Data curation, Writing – review & editing. Z. Chen: Formal analysis, Validation. B. Luo: Data curation, Validation. Z. Huang: Data curation, Visualization. X. Lin: Data curation, Validation. L. Xu: Data curation, Visualization. X. Li: Data curation. Y. Qu: Data curation. J. Xiao: Data curation. Z. Wang: Visualization. B. Shi: Data curation. Q.Ou: Visualization. B. Yao: Visualization. J. Yan: Data curation. Y. Zhuang: Data curation. Y. Zhang: Supervision, Validation, Writing – review & editing. R. Shi: Conceptualization, Formal analysis, Methodology, Supervision. Y. Xu: Investigation, Funding acquisition, Resources, Supervision, Project administration, Writing – review & editing.

---

**Extended Data Fig. 1.**
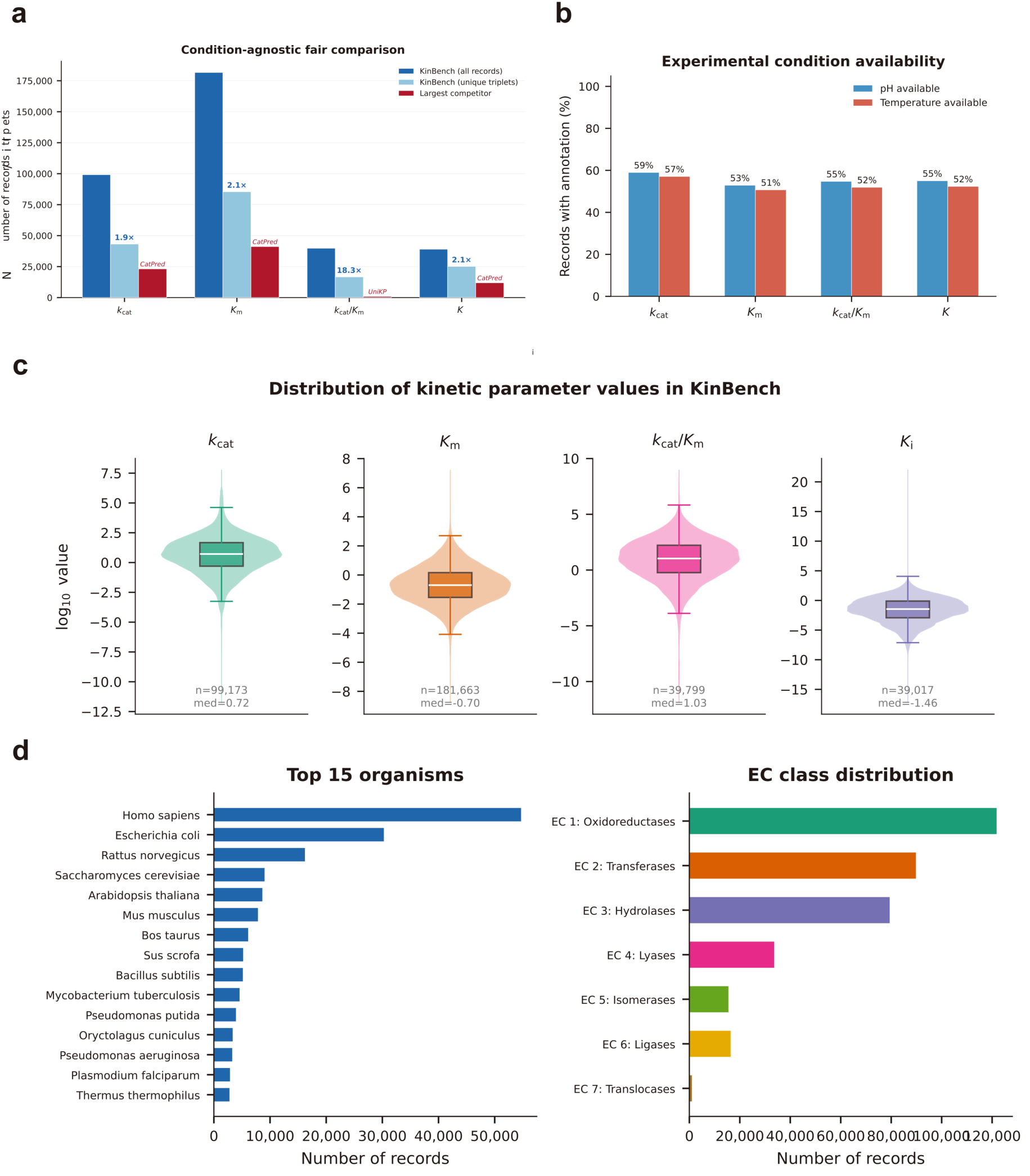
KinBench dataset characteristics and condition-agnostic validation. a, Condition-agnostic comparison: KinBench record counts, unique enzyme–substrate–organism triplets after geometric-mean aggregation of replicate measurements, and the largest task-specific competitor; KinBench retains 1.9× (*K*_*i*_ versus CatPred) to 18.3× (*k*_cat_/*K*_*m*_ versus UniKP) more entries per task under conservative accounting. b, Fraction of records with annotated pH and temperature per kinetic parameter; condition annotations provide training signal for EnzCast’s condition encoder. c, Distributions of log_10_-transformed kinetic values (violin and box plots); wide dynamic ranges justify log-space modelling. d, Top-15 organisms by record count and EC class distribution across all seven top-level enzyme classes; the benchmark spans 3,643 organisms and 4,442 EC classes.

**Extended Data Fig. 2.**
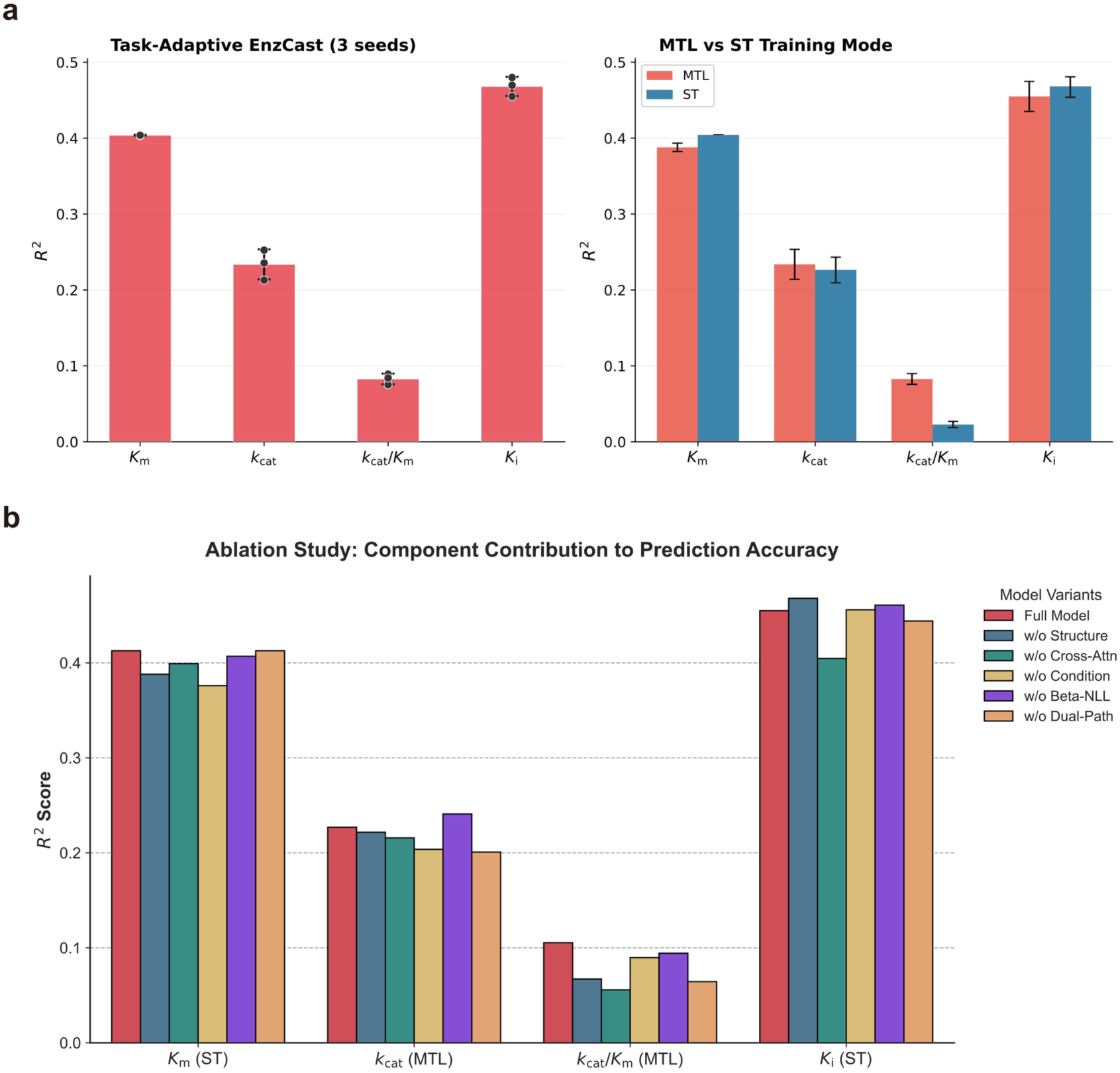
Multi-seed reproducibility and ablation study of EnzCast. a, Task-adaptive EnzCast *R*^2^ across three random seeds (42, 123, 2025). Bar heights indicate the mean; error bars indicate ± 1 s.d.; individual seed values are overlaid as points. Summary: *K*_*m*_, 0.404 ± 0.001 (CV 0.15%); *K*_*i*_, 0.468 ± 0.013 (CV 2.67%); *k*_cat_, 0.234 ± 0.020 (CV 8.47%); *k*_cat_/*K*_*m*_, 0.083 ± 0.007 (CV 8.45%). b, Comparison of multi-task learning (MTL) versus single-task (ST) training modes for each kinetic parameter; ST is preferred for *K*_*m*_and *K*_*i*_, and MTL for *k*_cat_ and *k*_cat_/*K*_*m*_due to the consistency constraint. c, Ablation study: grouped bar chart comparing *R*^2^of the full model against five ablation variants for each kinetic parameter under the task-adaptive training mode (ST for *K*_*m*_ and *K*_*i*_; MTL for *k*_cat_ and *k*_cat_/*K*_*m*_). Ablation variants remove: 3D structure encoder, substrate-to-structure cross-attention, experimental condition encoder, heteroscedastic *β*-NLL loss, and dual-pathway head. Removing any single component degrades performance on at least one task; cross-attention produces the largest average drop.

**Extended Data Fig. 3.**
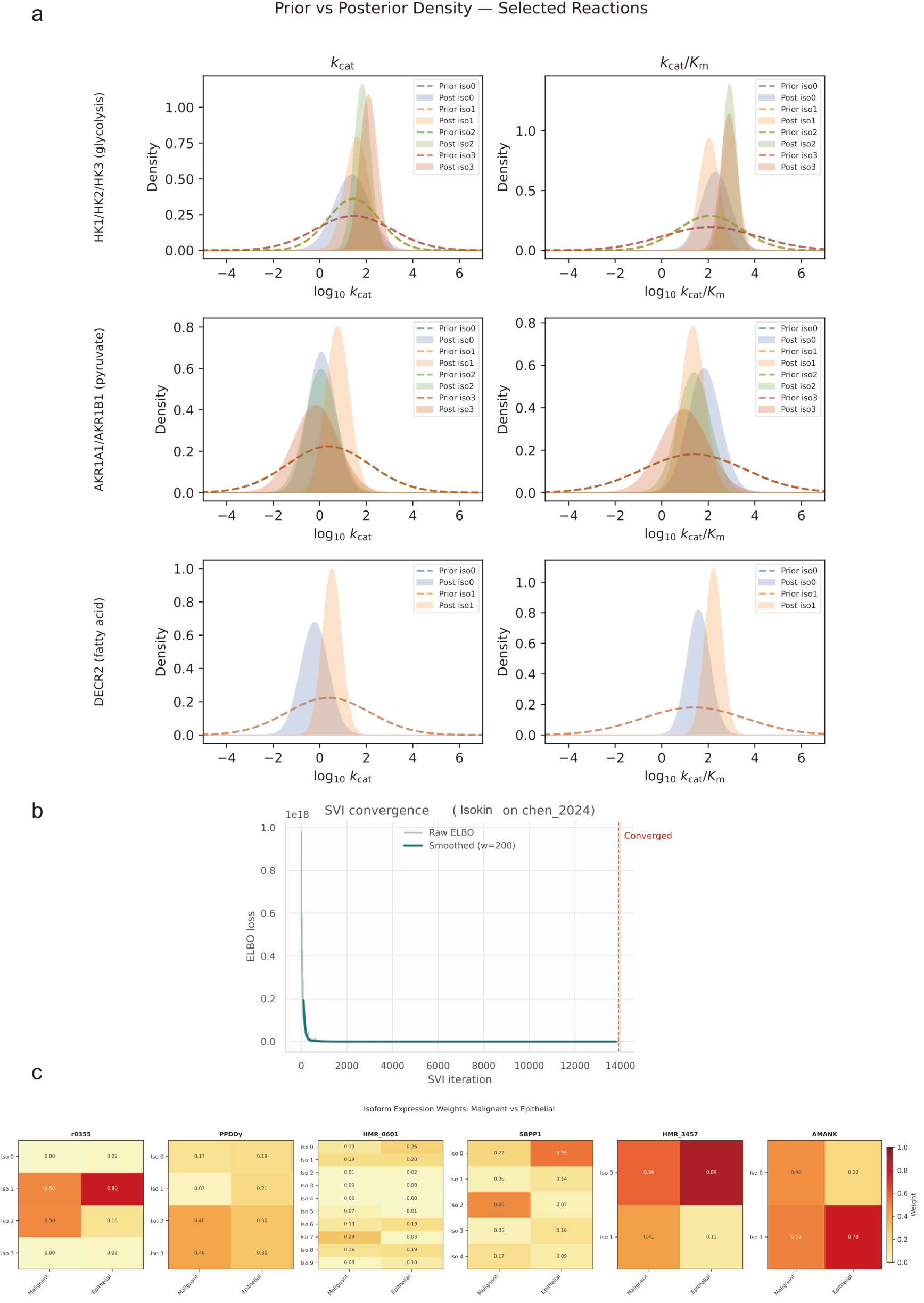
IsoKin framework, convergence diagnostics, and posterior characterization. a, Four-layer IsoKin schematic: isoform-level EnzCast prior assignment (pH 7.0, 37 °C); GPR decomposition via disjunctive normal form (OR branches: independent isoforms; AND clauses: rate-limiting complex aggregation); flux-constrained SVI with Recon3D stoichiometric constraints; expression-weighted effective kinetics derivation. b, SVI ELBO convergence on the CRC cohort [25]; early stopping at iteration 13,918 of 20,000. c, Prior versus posterior density overlays and isoform-weight heatmap for three representative reactions.

**Extended Data Fig. 4.**
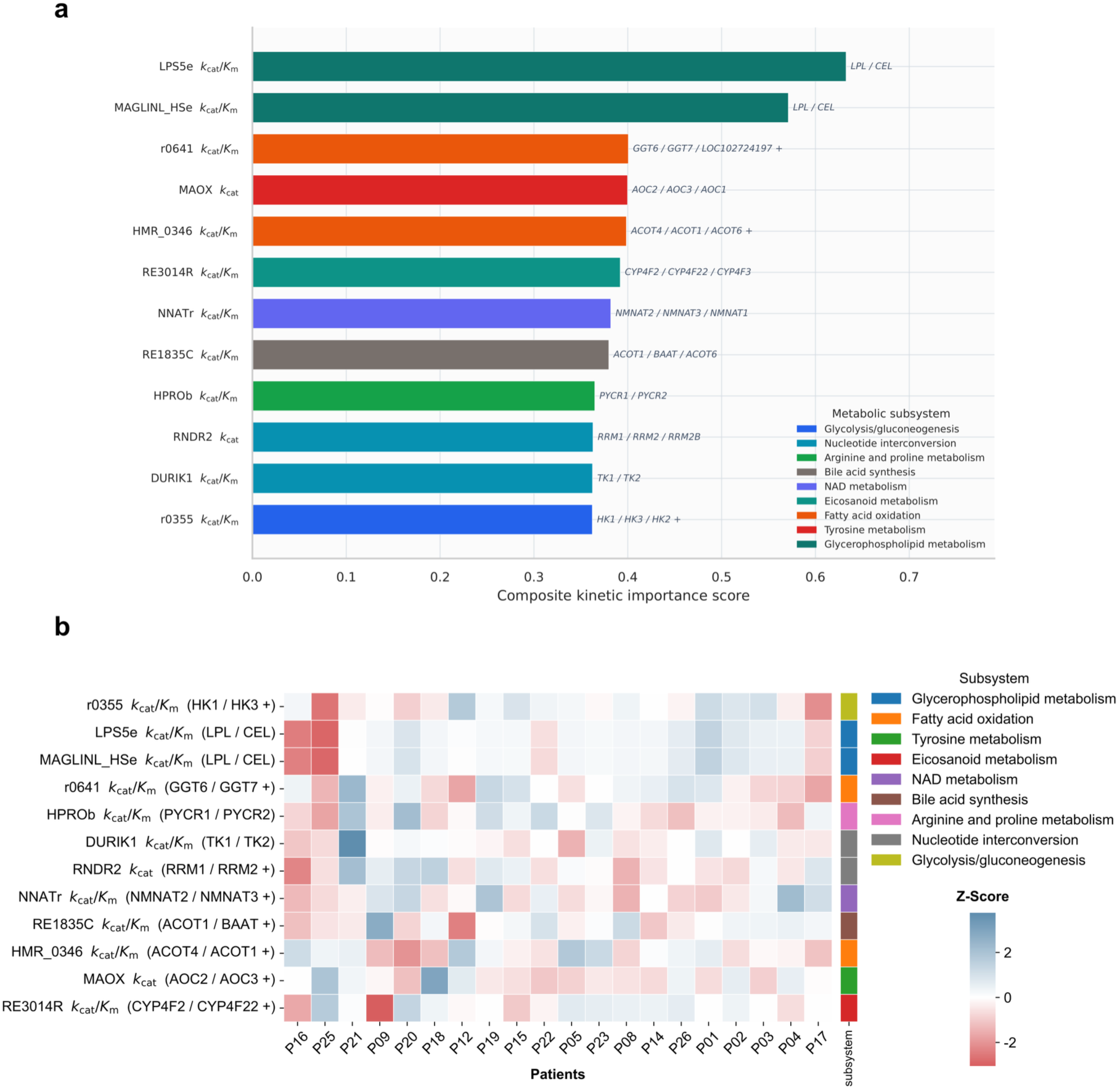
Composite reaction ranking and patient-by-reaction heatmap. a, Top 12 reactions by composite kinetic importance score in the CRC cohort [25]. Score integrates normalized kinetic shift score (weight 0.4), across-patient rewiring variability (weight 0.3), and mean posterior shrinkage (weight 0.3). Bar color encodes metabolic subsystem; gene family annotations are shown on each bar. b, Patient-by-reaction heatmap of malignant-vs.-epithelial rewiring z-scores for the 12 top-ranked interpretable reactions; left sidebar color-codes metabolic subsystem; columns (patients) and rows (reactions) are ordered by hierarchical clustering.

**Extended Data Fig. 5.**
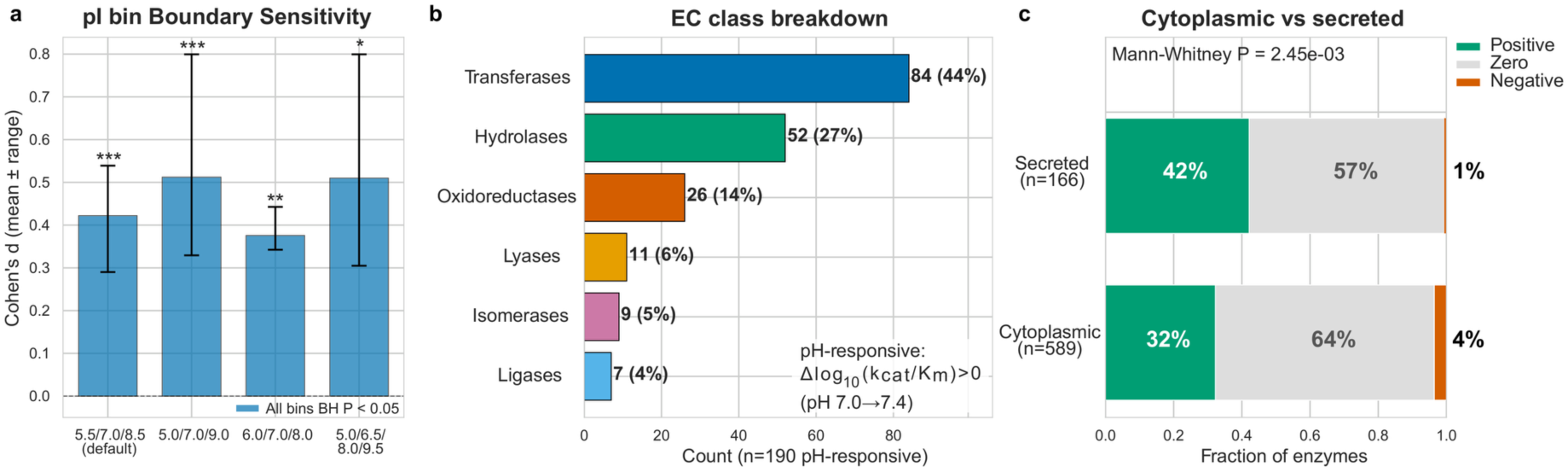
pH condition scan: pI sensitivity analysis and enzyme-class breakdown. a, Sensitivity of intracellular *k*_cat_/*K*_*m*_ shifts to pI bin boundaries; all individual bins remain significant (*P* < 0.05) across alternative binning schemes. b, Enzyme-class (EC top-level) breakdown of pH-responsive enzymes; transferases and hydrolases dominate among pH-responsive enzymes. c, Stacked fractions of positive, zero, and negative per-enzyme *Δ*log_10_(*k*_cat_/*K*_*m*_) responses for secreted (top) and cytoplasmic (bottom) enzymes under intracellular alkalinization (Mann–Whitney *P* = 2.5 × 10^−3^).

**Extended Data Fig. 6.**
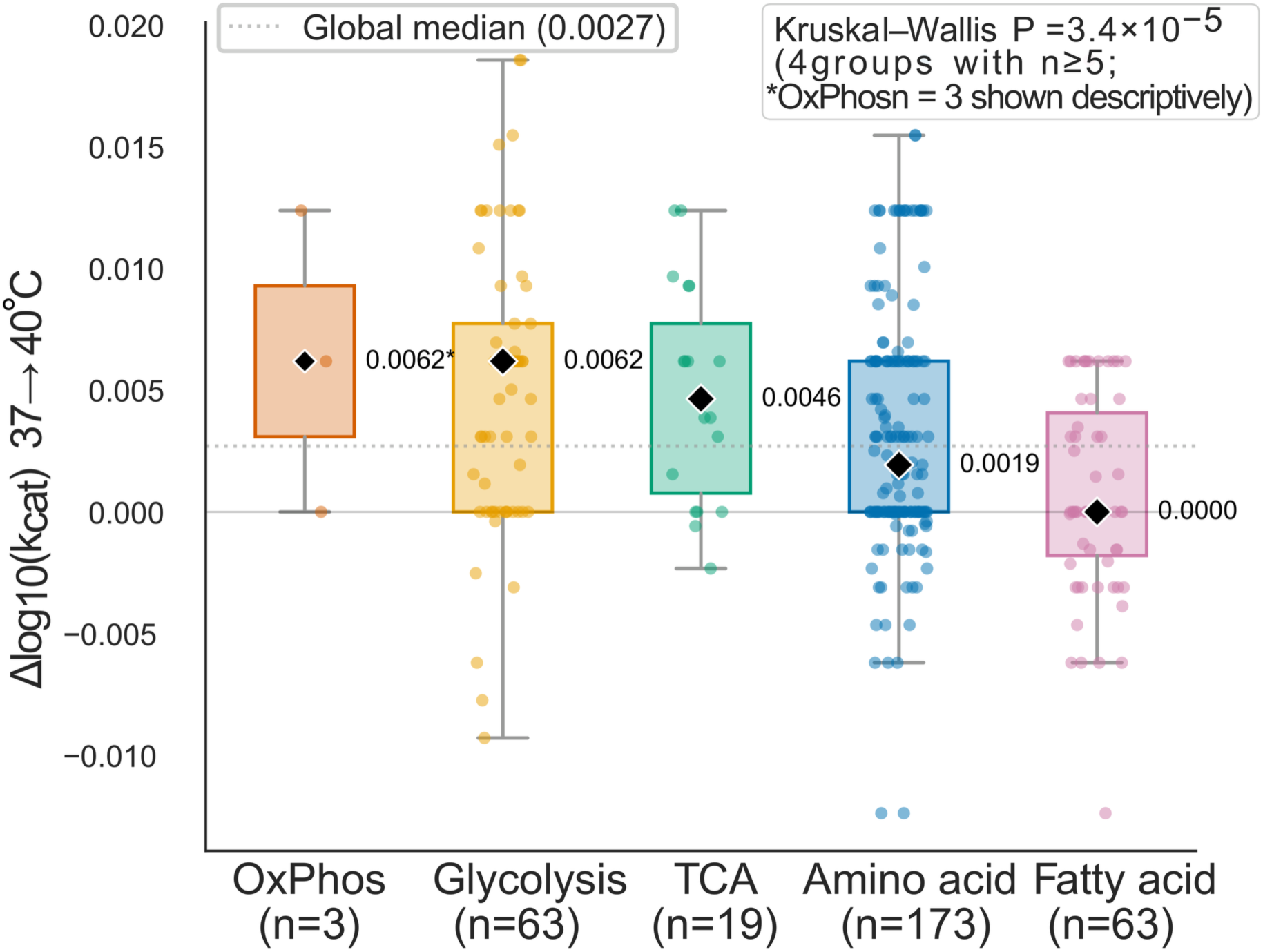
Temperature condition scan: per-subsystem metabolic pathway response. Distribution of *Δ*log_10_(*k*_cat_) from 37 to 40 °C across metabolic subsystems, with median values indicated; oxidative phosphorylation, glycolysis, and TCA cycle subsystems are highlighted.

**Extended Data Fig.7.**
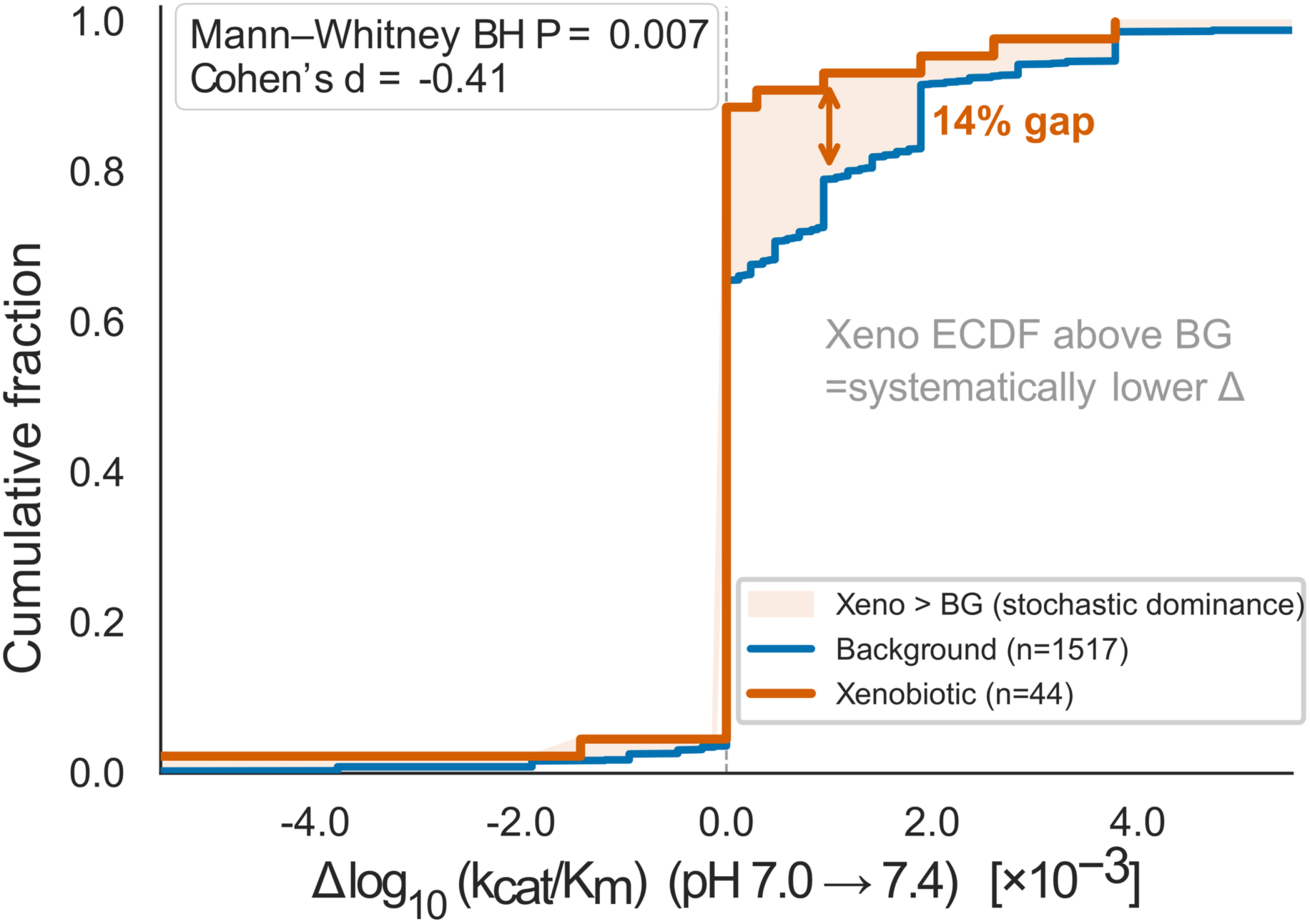
Xenobiotic-metabolism enzyme kinetics under intracellular alkalinization. Empirical cumulative distribution functions (ECDFs) of *Δ*log_10_(*k*_cat_/*K*_*m*_) under pH 7.0 → 7.4 for 44 xenobiotic-metabolism enzymes (red) versus the metabolic background (*n* = 1,517; blue). The xenobiotic ECDF sits consistently above the background curve, indicating that xenobiotic enzymes exhibit systematically lower catalytic-efficiency gains under intracellular alkalinization (Mann–Whitney BH *P* = 0.007, Cohen’s *d* = −0.41).

**Extended Data Fig. 8.**
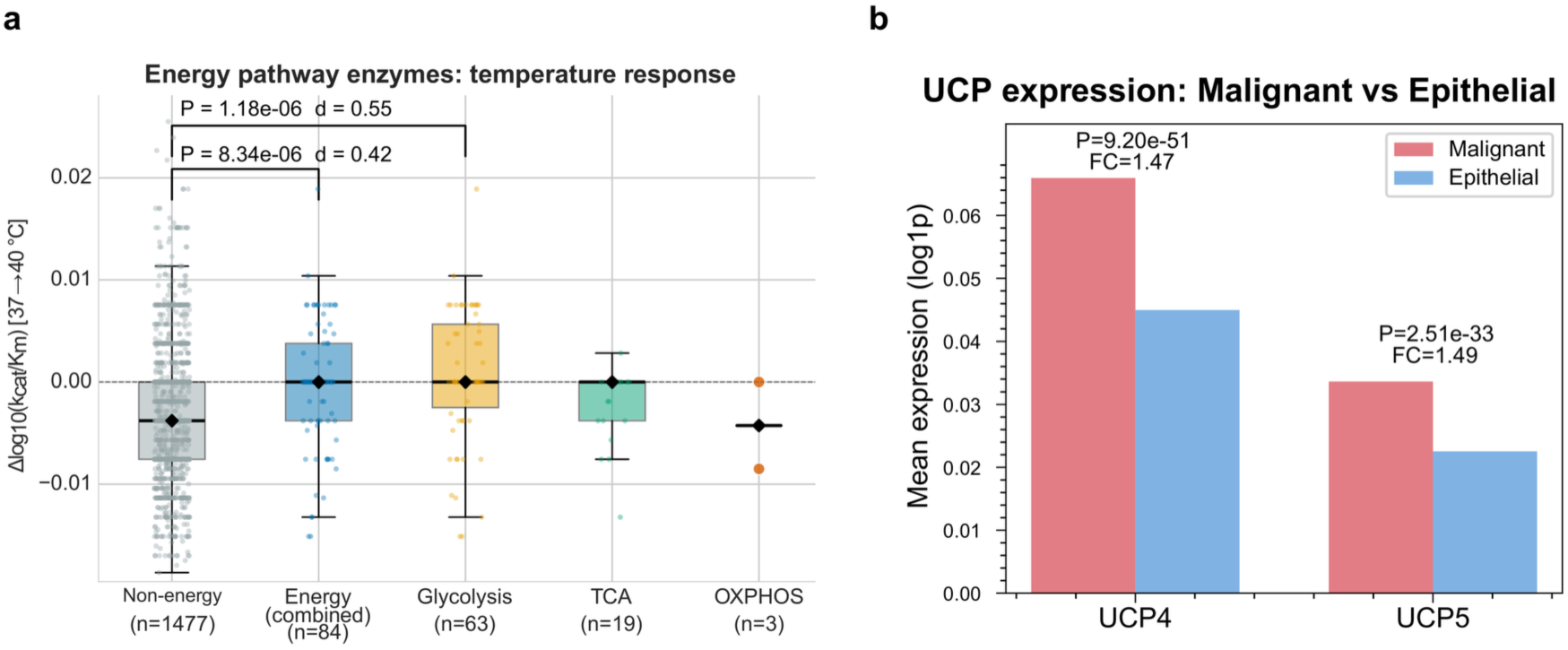
Energy-production pathway temperature sensitivity and mitochondrial uncoupling protein expression. a, Boxplots comparing *Δ*log_10_(*k*_cat_/*K*_*m*_) under 37 → 40 °C for energy-production pathway enzymes (glycolysis, TCA, OXPHOS; *n* = 84) versus non-energy background (Mann–Whitney *P* = 8.3 × 10^−6^, *d* = 0.42); glycolytic enzymes (*n* = 63) drive the strongest signal (*P* = 1.18 × 10^−6^, *d* = 0.55). b, Single-cell expression of mitochondrial uncoupling proteins UCP4 (SLC25A27; fold change = 1.47, *P* = 9.2 × 10^−51^) and UCP5 (SLC25A14; fold change = 1.49, *P* = 2.5 × 10^−33^) in malignant versus epithelial cells from the CRC cohort.

**Extended Data Fig. 9.**
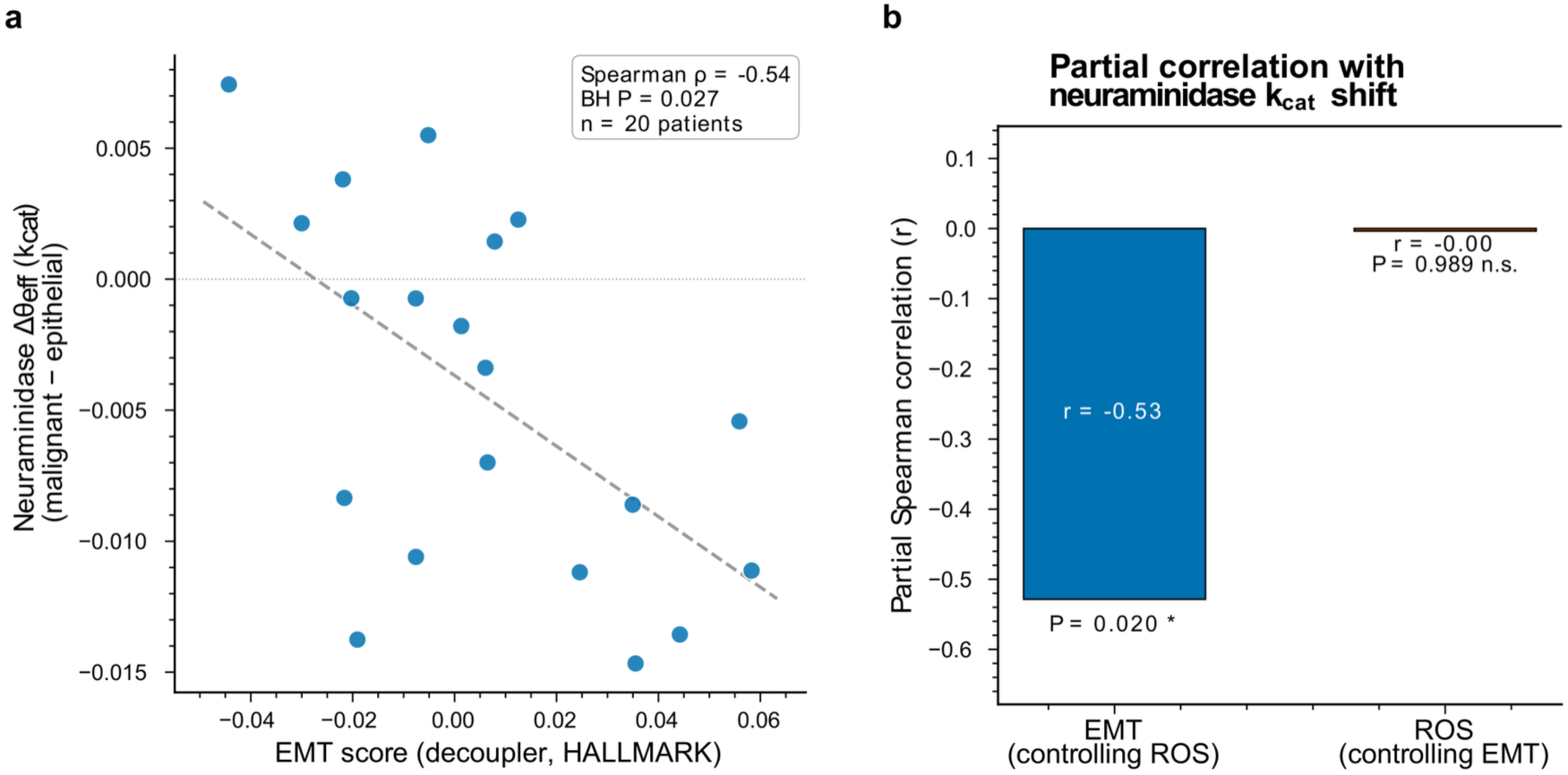
Neuraminidase kinetics and epithelial–mesenchymal transition in the CRC cohort. a, Patient-level scatter plot of EMT score versus signed malignant-vs.-epithelial neuraminidase *k*_cat_ shift (Spearman *ρ* = −0.54, BH *P* = 0.027). b, Partial correlation bar chart: the neuraminidase–EMT association is independent of oxidative stress (ROS): partial *r* = −0.53, *P* = 0.020 after controlling for ROS, whereas the ROS contribution is negligible (*r* = −0.003, *P* = 0.989).

## Appendix A Extended Data

**Extended Data Table 1.**
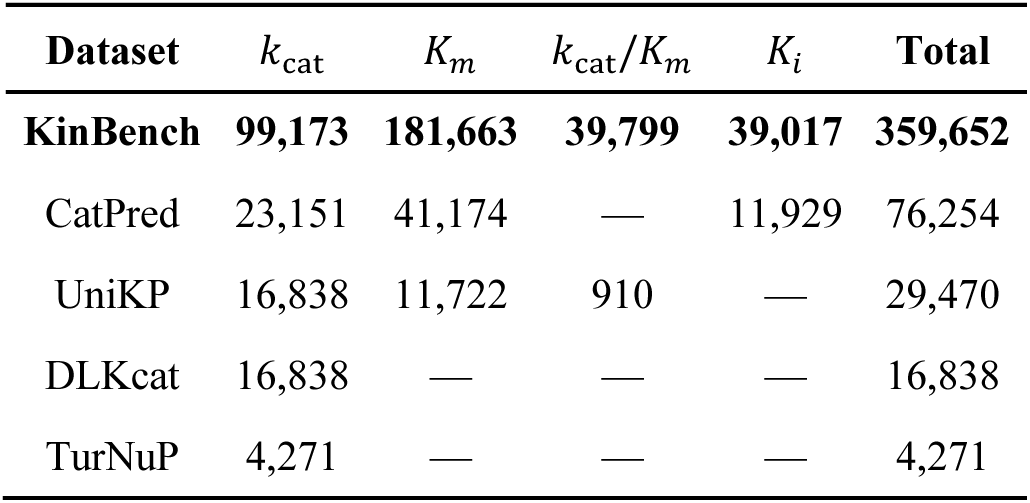
Comparison of KinBench with existing enzyme kinetic ML benchmarks. Record counts per kinetic parameter for each benchmark definition used in the Section 2.1 comparison. A dash (—) indicates that the parameter is not covered. KinBench is the only benchmark covering all four parameters and is the largest for each directly comparable task.

| Dataset | $k_{\text{cat}}$ | $K_m$ | $k_{\text{cat}}/K_m$ | $K_i$ | Total |
| --- | --- | --- | --- | --- | --- |
| **KinBench** | **99,173** | **181,663** | **39,799** | **39,017** | **359,652** |
| CatPred | 23,151 | 41,174 | — | 11,929 | 76,254 |
| UniKP | 16,838 | 11,722 | 910 | — | 29,470 |
| DLKcat | 16,838 | — | — | — | 16,838 |
| TurNuP | 4,271 | — | — | — | 4,271 |

**Extended Data Table 2.** Full benchmark metrics for all methods on the KinBench UniRef50-clustered test set. EnzCast-ST and EnzCast-MTL denote single-task and multi-task training modes, respectively. The task-adaptive strategy (bold) selects ST for *K*_*m*_ and *K*_*i*_, MTL for *k*_cat_ and *k*_cat_/*K*_*m*_.

| Method | Task | $R^2$ | RMSE | MAE | Pearson | Spearman |
| --- | --- | --- | --- | --- | --- | --- |
| **EnzCast-ST** | $K_m$ | **0.413** | **0.959** | **0.730** | **0.645** | **0.648** |
| EnzCast-MTL | $K_m$ | 0.368 | 0.995 | 0.756 | 0.611 | 0.620 |
| UniKP | $K_m$ | 0.401 | 0.969 | 0.741 | 0.639 | 0.640 |
| TurNuP | $K_m$ | 0.365 | 0.998 | 0.773 | 0.623 | 0.625 |
| CatPred | $K_m$ | 0.293 | 1.053 | 0.800 | 0.555 | 0.563 |
| DLKcat | $K_m$ | 0.183 | 1.132 | 0.873 | 0.488 | 0.493 |
| **EnzCast-MTL** | $k_{\text{cat}}$ | **0.227** | **1.373** | **1.062** | **0.481** | **0.466** |
| EnzCast-ST | $k_{\text{cat}}$ | 0.220 | 1.379 | 1.059 | 0.481 | 0.456 |
| UniKP | $k_{\text{cat}}$ | 0.192 | 1.404 | 1.088 | 0.474 | 0.442 |
| TurNuP | $k_{\text{cat}}$ | 0.178 | 1.416 | 1.095 | 0.437 | 0.413 |
| CatPred | $k_{\text{cat}}$ | 0.125 | 1.461 | 1.122 | 0.383 | 0.374 |
| DLKcat | $k_{\text{cat}}$ | -0.174 | 1.691 | 1.317 | 0.167 | 0.145 |
| **EnzCast-MTL** | $k_{\text{cat}}/K_m$ | **0.105** | **1.993** | **1.456** | **0.334** | **0.388** |
| EnzCast-ST | $k_{\text{cat}}/K_m$ | 0.022 | 2.084 | 1.480 | 0.200 | 0.288 |
| UniKP | $k_{\text{cat}}/K_m$ | 0.072 | 2.029 | 1.490 | 0.274 | 0.311 |
| TurNuP | $k_{\text{cat}}/K_m$ | 0.061 | 2.041 | 1.503 | 0.250 | 0.274 |
| CatPred | $k_{\text{cat}}/K_m$ | -0.037 | 2.145 | 1.625 | 0.207 | 0.227 |
| DLKcat | $k_{\text{cat}}/K_m$ | -0.167 | 2.275 | 1.719 | 0.126 | 0.095 |
| **EnzCast-ST** | $K_i$ | **0.455** | **1.570** | **1.177** | **0.676** | **0.688** |
| EnzCast-MTL | $K_i$ | 0.444 | 1.586 | 1.165 | 0.669 | 0.687 |
| UniKP | $K_i$ | 0.460 | 1.562 | 1.183 | 0.696 | 0.702 |
| TurNuP | $K_i$ | 0.413 | 1.629 | 1.220 | 0.658 | 0.700 |
| CatPred | $K_i$ | 0.354 | 1.709 | 1.298 | 0.602 | 0.633 |
| DLKcat | $K_i$ | 0.280 | 1.804 | 1.369 | 0.547 | 0.560 |

**Extended Data Table 3.** Ablation study results. *R*^2^ on the KinBench UniRef50-clustered test set when each architectural component is removed. Task-adaptive mode is used: MTL for *k*_cat_ and *k*_cat_/*K*_*m*_, ST for *K*_*m*_ and *K*_*i*_. Removing any single component degrades performance on at least one task.

| Configuration | $K_m$ (ST) | $k_{\text{cat}}$ (MTL) | $k_{\text{cat}}/K_m$ (MTL) | $K_i$ (ST) |
| --- | --- | --- | --- | --- |
| Full model | 0.413 | 0.227 | 0.105 | 0.455 |
| w/o Structure encoder | 0.388 | 0.222 | 0.067 | 0.468 |
| w/o Cross-attention | 0.399 | 0.216 | 0.056 | 0.405 |
| w/o Condition encoder | 0.376 | 0.204 | 0.090 | 0.456 |
| w/o β-NLL loss | 0.407 | 0.241 | 0.094 | 0.461 |
| w/o Dual-pathway head | 0.413 | 0.201 | 0.064 | 0.444 |

**Extended Data Table 4.** BH-significant kinetic differences across malignant-vs.-non-malignant cell-type comparisons in the CRC cohort. For each of the 1,842 IsoKin-resolved reactions and each catalytic parameter, the table reports the number and percentage of reactions whose malignant-vs.-non-malignant posterior shift survives Benjamini–Hochberg adjustment ( **q** < 0.05). The malignant-vs.-originating-epithelial baseline yields an approximately threefold (2.9–3.2×) smaller differential than malignant-vs.-stromal comparisons, indicating that cell-type identity contributes substantially to the apparent kinetic divergence.

| Comparison | $k_{\text{cat}}$ BH-significant (n, %) | $k_{\text{cat}}/K_m$ BH-significant (n, %) |
| --- | --- | --- |
| Malignant vs. Macrophage | 705 / 1,842 (38.3%) | 744 / 1,842 (40.4%) |
| Malignant vs. Fibroblast | 716 / 1,842 (38.9%) | 714 / 1,842 (38.8%) |
| Malignant vs. Originating Epithelial | 245 / 1,842 (13.3%) | 233 / 1,842 (12.6%) |

### Implementation fidelity verification

To ensure that performance differences observed on KinBench reflect genuine model capability rather than implementation artifacts, we retrained each baseline on its original benchmark dataset and compared with published metrics (Extended Data Table 5). UniKP achieved the closest reproduction across all baselines, with reported *R*^2^values for *K*_*m*_and *k*_cat_/*K*_*m*_reproduced within 0.03–0.06. For *k*_cat_, where the original publication reports PCC rather than *R*^2^, our Pearson correlation (0.81) closely matched the reported 0.85, supporting functional alignment of the ProtT5 + SMILES Transformer + ExtraTreesRegressor pipeline with the original implementation. CatPred was evaluated in simplified mode (single model, MSE loss) rather than the original 10-model ensemble with MVE loss. Despite this simplification, all three tasks maintained *R*^2^ within 0.03–0.10 of the reported ensemble values (*k*_cat_: 0.55 vs 0.60; *K*_*m*_: 0.54 vs 0.64; *K*_*i*_: 0.60 vs 0.63). All three tasks maintained strong correlations (Pearson *r* = 0.74 − 0.78). DLKcat showed the largest *R*^2^ gap relative to other baselines (*R*^2^ = 0.49). The original DLKcat paper does not report *R*^2^; it reports PCC = 0.71 and RMSE = 1.06 on the test set. Our PCC of 0.71 exactly matches the reported value, confirming architectural correctness. TurNuP *k*_cat_ was reproduced using 10-fold cross-validation matching the original protocol, with ESM-1b embeddings re-extracted from protein sequences and structural fingerprints parsed from the original dataset. Our reproduction achieved *R*^2^ = 0.637 (Pearson *r* = 0.81), substantially exceeding the originally reported *R*^2^of ∼0.44. The improvement is likely due in part to re-extracting full ESM-1b embeddings rather than using the pre-computed embeddings in the source CSV, which appeared truncated. TurNuP *K*_*m*_reproduction was not attempted because 92.6% of target values in the available dataset are null. Across all reproductions, the consistently high Pearson correlation coefficients (*r* ≥ 0.70) and Spearman rank correlations (*ρ* ≥ 0.69) demonstrate that our reimplementations faithfully capture each baseline’s learned feature-to-target mappings. The *R*^2^gaps observed for DLKcat and CatPred *k*_cat_are attributable to known, quantifiable differences in training configuration rather than architectural errors, and do not compromise the fairness of the KinBench comparison.

**Extended Data Table 5.** Baseline implementation fidelity verification. Each baseline was retrained on its original benchmark dataset using our reimplementation. All metrics are computed in log_10_space. “Reported” values are from the respective original publications. PCC: Pearson correlation coefficient.

| Baseline | Task | $N_{\text{test}}$ | Split | $R^2$ (ours) | $R^2$ (rep.) | RMSE | MAE | PCC (ours) | Status |
| --- | --- | --- | --- | --- | --- | --- | --- | --- | --- |
| DLKcat | $k_{\text{cat}}$ | 1,701 | 80/10/10 | 0.494 | —<sup>†</sup> | 1.062 | 0.779 | 0.709 | Notea |
| UniKP | $k_{\text{cat}}$ | 3,401 | 80/20 | 0.654 | —<sup>†</sup> | 0.873 | 0.588 | 0.809 | ✓ |
| UniKP | $K_m$ | 2,345 | 80/20 | 0.499 | 0.530 | 0.843 | 0.637 | 0.709 | ✓ |
| UniKP | $k_{\text{cat}}/K_m$ | 914 | 5-fold CV | 0.594 | 0.65 | 1.163 | 0.823 | 0.771 | ✓ |
| TurNuP | $k_{\text{cat}}$ | 4,271 | 10-fold CV | 0.637 | 0.44 | 0.715 | 0.528 | 0.809 | Notec |
| CatPred | $k_{\text{cat}}$ | 2,316 | 80/10/10 | 0.546 | 0.602 | 1.109 | 0.760 | 0.749 | Noteb |
| CatPred | $K_m$ | 4,110 | 80/10/10 | 0.542 | 0.642 | 0.882 | 0.661 | 0.742 | Noteb |
| CatPred | $K_i$ | 1,191 | 80/10/10 | 0.601 | 0.632 | 1.246 | 0.888 | 0.784 | Noteb |

<sup>†</sup>DLKcat and UniKP ( $k_{\text{cat}}$ ) do not report $R^2$ in their original publications; they report PCC = 0.71 and PCC = 0.85, respectively.
<sup>a</sup>Our reimplementaion uses validation-based model selection (80/10/10 split). The original DLKcat reports results on an 80/20 split without explicit early-stopping details. The $R^2$ gap may reflect undocumented training regularisation; the PCC of 0.71 confirms correct predictive behaviour.
<sup>b</sup>Simplified mode (single model, MSE loss). The original CatPred uses a 10-model ensemble with MVE loss, which typically improves $R^2$ by 0.05–0.10. Reported $R^2$ values are from CatPred Table 2 (held-out test, ensemble).
<sup>c</sup>Our reproduction substantially exceeds the originally reported $R^2$ (0.637 vs 0.44), primarily because ESM-1b embeddings were re-extracted from full protein sequences rather than using the pre-computed embeddings in the TurNuP dataset, which appeared truncated.
UniKP $k_{\text{cat}}/K_m$ : 914 valid samples after filtering 231 unparseable SMILES (20.2%) from the original 1,145-sample dataset; confirmed as an upstream data issue. TurNuP $K_m$ reproduction was not attempted (92.6% null targets in source data).

## REFERENCES

[1] Bar-Even, A. et al. The moderately efficient enzyme: Evolutionary and physicochemical trends shaping enzyme parameters. Biochemistry 50, 4402–4410 (2011).

[2] Davidi, D. et al. Global characterization of in vivo enzyme catalytic rates and their correspondence to in vitro kcat measurements. Proceedings of the National Academy of Sciences 113, 3401–3406 (2016).

[3] Bisswanger, H. Enzyme assays. Perspectives in Science 1, 41–55 (2014).

[4] Webb, B. A., Chimenti, M., Jacobson, M. P. & Bhargava, R. Dysregulated pH: a perfect storm for cancer progression. Nature Reviews Cancer 11, 671–677 (2011).

[5] Corbet, C. & Feron, O. Tumour acidosis: from the passenger to the driver’s seat. Nature Reviews Cancer 17, 577–593 (2017).

[6] Repasky, E. A., Evans, S. S. & Dewhirst, M. W. Temperature matters! and why it should matter to tumor immunologists. Cancer Immunology Research 1, 210–216 (2013).

[7] Daniel, R. M. & Danson, M. J. The role of dynamics in enzyme activity. FEBS Journal 280, 5981–5990 (2013).

[8] Chio, I. I. C. & Tuveson, D. A. ROS in cancer: The burning question. Trends in Molecular Medicine 23, 411–429 (2017).

[9] Cantó, C., Menzies, K. J. & Auwerx, J. NAD+ metabolism and the control of energy homeostasis: A balancing act between mitochondria and the nucleus. Cell Metabolism 22, 31–53 (2015).

[10] Li, F. et al. Deep learning-based kcat prediction enables improved enzyme-constrained model reconstruction. Nature Catalysis 5, 662–672 (2022).

[11] Yu, H., Deng, H., He, J., Keasling, J. D. & Luo, X. UniKP: a unified framework for the prediction of enzyme kinetic parameters. Nature Communications 14, 8211 (2023).

[12] Kroll, A., Rousset, Y., Hu, X.-P., Liebrand, N. A. & Lercher, M. J. Turnover number predictions for kinetically uncharacterized enzymes using machine and deep learning. Nature Communications 14, 4139 (2023).

[13] Boorla, V. S. & Maranas, C. D. CatPred: a comprehensive framework for deep learning in vitro enzyme kinetic parameters. Nature Communications 16, 2072 (2025).

[14] Lin, Z. et al. Evolutionary-scale prediction of atomic-level protein structure with a language model. Science 379, 1123–1130 (2023). ESMFold.

[15] Hu, E. J., et al. LoRA: Low-rank adaptation of large language models. International Conference on Learning Representations (2022).

[16] Satorras, V. G., Hoogeboom, E. & Welling, M. E(n) equivariant graph neural networks. International Conference on Machine Learning 9323–9332 (2021).

[17] Xu, K., Hu, W., Leskovec, J. & Jegelka, S. How powerful are graph neural networks? International Conference on Learning Representations (2019).

[18] Rogers, D. & Hahn, M. Extended-connectivity fingerprints. Journal of Chemical Information and Modeling 50, 742–754 (2010).

[19] Steinegger, M. & Söding, J. MMseqs2 enables sensitive protein sequence searching for the analysis of massive data sets. Nature Biotechnology 35, 1026–1028 (2017).

[20] Brunk, E. et al. Recon3D enables a three-dimensional view of gene variation in human metabolism. Nature Biotechnology 36, 272–281 (2018).

[21] Hoffman, M. D., Blei, D. M., Wang, C. & Paisley, J. Stochastic variational inference. Journal of Machine Learning Research 14, 1303–1347 (2013).

[22] Wittig, U. et al. SABIO-RK—database for biochemical reaction kinetics. Nucleic Acids Research 40, D742–D748 (2012).

[23] Chang, A. et al. BRENDA, the ELIXIR core data resource in 2021: new developments and updates. Nucleic Acids Research 49, D498–D508 (2021).

[24] The UniProt Consortium. UniProt: the universal protein knowledgebase in 2023. Nucleic Acids Research 51, D523–D531 (2023).

[25] Chen, B., Li, M. et al. Single-cell and spatial transcriptomics reveal a high-resolution view of the tumor microenvironment in colorectal cancer. Cancer Cell 42, 1726–1744 (2024).

[26] Alghamdi, N. et al. A graph neural network model to estimate cell-wise metabolic flux using single-cell RNA-seq data. Genome Research 31, 1867–1884 (2021).

[27] Tsherniak, A. et al. Defining a cancer dependency map. Cell 170, 564–576 (2017).

[28] The Cancer Genome Atlas Network. Comprehensive molecular characterization of human colon and rectal cancer. Nature 487, 330–337 (2012).

29. Landrum, G. RDKit: Open-source cheminformatics. https://www.rdkit.org (2016). Software available at https://www.rdkit.org.

[30] Berman, H. M. et al. The Protein Data Bank. Nucleic Acids Research 28, 235–242 (2000).

[31] Jumper, J. et al. Highly accurate protein structure prediction with AlphaFold. Nature 596, 583–589 (2021).

[32] Gilmer, J., Schoenholz, S. S., Riley, P. F., Vinyals, O. & Dahl, G. E. Neural message passing for quantum chemistry. Proceedings of the 34th International Conference on Machine Learning 70, 1263–1272 (2017).

[33] Seitzer, M., Tavakoli, A., Antic, D. & Martius, G. On the pitfalls of heteroscedastic uncertainty estimation with probabilistic neural networks. International Conference on Learning Representations (2022).

[34] Kendall, A., Gal, Y. & Cipolla, R. Multi-task learning using uncertainty to weigh losses for scene geometry and semantics. Proceedings of the IEEE Conference on Computer Vision and Pattern Recognition 7482–7491 (2018).

[35] Chen, T., Kornblith, S., Norouzi, M. & Hinton, G. A simple framework for contrastive learning of visual representations. International Conference on Machine Learning 1597–1607 (2020).

[36] Loshchilov, I. & Hutter, F. SGDR: stochastic gradient descent with warm restarts. Proceedings of the 5th International Conference on Learning Representations (2017).

[37] Bingham, E. et al. Pyro: Deep universal probabilistic programming. Journal of Machine Learning Research 20, 1–6 (2019).

[38] Pavlova, N. N. & Thompson, C. B. The emerging hallmarks of cancer metabolism. Cell Metabolism 23, 27–47 (2016).

[39] Wolf, F. A., Angerer, P. & Theis, F. J. SCANPY: large-scale single-cell gene expression data analysis. Genome Biology 19, 15 (2018).

[40] Wolock, S. L., Lopez, R. & Klein, A. M. Scrublet: Computational identification of cell doublets in single-cell transcriptomic data. Cell Systems 8, 281–291 (2019).

[41] Korsunsky, I. et al. Fast, sensitive and accurate integration of single-cell data with Harmony. Nature Methods 16, 1289–1296 (2019).

[42] Traag, V. A., Waltman, L. & van Eck, N. J. From Louvain to Leiden: guaranteeing well-connected communities. Scientific Reports 9, 5233 (2019).

[43] Elnaggar, A. et al. ProtTrans: Toward understanding the language of life through self-supervised learning. IEEE Transactions on Pattern Analysis and Machine Intelligence 44, 7112–7127 (2022).

[44] Benjamini, Y. & Hochberg, Y. Controlling the false discovery rate: A practical and powerful approach to multiple testing. Journal of the Royal Statistical Society: Series B 57, 289–300 (1995).

